# The robustness of a simple dynamic model of island biodiversity to geological and eustatic change

**DOI:** 10.1101/2021.07.26.453064

**Authors:** Pedro Santos Neves, Joshua W. Lambert, Luis Valente, Rampal S. Etienne

**Affiliations:** Groningen Institute for Evolutionary Life Sciences, University of Groningen, Box 11103, 9700 CC Groningen, The Netherlands.; Naturalis Biodiversity Center, Darwinweg 2, 2333 CR Leiden, The Netherlands

**Author notes:** These authors contributed equally to this work (Pedro Santos Neves, Joshua W. Lambert). Corresponding Author: Pedro Santos Neves, Groningen Institute for Evolutionary Life Sciences, Box 11103, 9700 CC Groningen, The Netherlands.

## Abstract

**Aim:** Biodiversity on islands is affected by various geo-physical processes and sea-level fluctuations. Oceanic islands (never connected to a landmass) are initially vacant with diversity accumulating via colonisation and speciation, followed by a decline as islands shrink. Continental islands have species upon formation (when disconnected from the mainland) and may have transient land-bridge connections. Theoretical predictions for the effects of these geo-processes on rates of colonisation, speciation and extinction have been proposed, but methods of phylogenetic inference assume only oceanic island scenarios without accounting for island ontogeny, sea-level changes or past landmass connections. Here, we analyse to what extent ignoring geodynamics affects the inference performance of a phylogenetic island model, DAISIE, when confronted with simulated data that violate its assumptions.

**Location:** Simulation of oceanic and continental islands.

**Methods:** We extend the DAISIE simulation model to include: area-dependent rates of colonisation and diversification associated with island ontogeny and sea-level fluctuations, and continental islands with biota present upon separation from the mainland, and shifts in rates to mimic temporary land-bridges. We quantify the error made when geo-processes are not accounted for by applying DAISIE’s inference method to geodynamic simulations.

**Results:** We find that the robustness of the model to dynamic island area is high (error is small) for oceanic islands and for continental islands that have been separated for a long time, suggesting that, for these island types, it is possible to obtain reliable results when ignoring geodynamics. However, for continental islands that have been recently or frequently connected, robustness of DAISIE is low, and inference results should not be trusted.

Main conclusions: This study highlights that under a large proportion of island biogeographic geo-scenarios (oceanic islands and ancient continental fragments) a simple phylogenetic model ignoring geodynamics is empirically applicable and informative. However, recent connection to the continent cannot be ignored, requiring development of a new inference model.

## Introduction

The study of biodiversity on islands rests upon the foundational work of MacArthur and Wilson’s (1963; 1967) equilibrium theory of biodiversity (ETIB). The theory describes island diversity as determined by species colonisation and extinction, governed by island area and isolation. The ETIB proposes that an equilibrium state of biodiversity emerges when the rates of colonisation and extinction are equal. Additionally, diversity can accumulate through *in situ* speciation, particularly on large isolated islands (MacArthur and Wilson, 1967; Losos and Schluter, 2000; Rosindell and Phillimore, 2011; Valente et al., 2020). While the ETIB has found empirical support at both ecological (Simberloff and Wilson, 1970) and evolutionary (Valente et al., 2017b, 2020) time scales and while its fundamental mechanisms are generally accepted, the state of equilibrium is thought to not be easily reached or maintained (Heaney, 2000; Whittaker et al., 2008; Valente et al., 2014; Warren et al., 2015; Fernández-Palacios et al., 2016; Marshall and Quental, 2016), because of frequent ecological, geological and eustatic perturbations. The general dynamic model (GDM), glacial sensitive model (GSM), and sea-level sensitive model (SLS) of island biogeography extend the ETIB by incorporating the influence of geodynamics (island ontogeny and sea-level) on evolutionary processes (Whittaker et al., 2008; Fernández-Palacios et al., 2016; Ávila et al., 2019). Modelling the GDM and GSM has revealed empirical support for an effect of geodynamics on biodiversity (Whittaker et al., 2008; Bunnefeld and Phillimore, 2012; Steinbauer et al., 2013; Rijsdijk et al., 2014; Lim and Marshall, 2017).

The trajectory of island diversity is a result of ecological and geological evolution. Among the relevant physical and biological island characteristics that influence island diversity, area stands out (Ali, 2017). Islands are classified as oceanic or continental, differentiated by the latter having had a physical connection to a mainland at some point in their history (Wallace, 1880; Ali, 2018). Continental islands form via a separation from a continental landmass and are stable or undergo a very slow decline in island area (Ali, 2018). Here we use these broad terms but acknowledge the pronounced geological differences between and within various island types (see Ali (2017, 2018)). Island ontogeny encapsulates the geological evolution of area, topographic complexity, and elevation over an island’s lifespan, which impact species diversity. Oceanic islands generally undergo an ontogeny starting with initial island emergence, an uplift phase of rapid increase in area, until the island reaches a maximum size. This is followed by a relatively slow decline in area, and eventually atoll formation and submergence (Ramalho et al., 2013; Ali, 2017). Oceanic island ontogeny is hypothesised to affect diversity. For example, the GDM uses a niche-based argument for a time-variable carrying capacity that governs speciation, extinction and colonisation as area changes (Whittaker et al., 2008). Simulations of the effects of island ontogeny on phylogenetic data show that island ontogeny can indeed influence species richness, but its effect depends on specific characteristics of the island in question (Valente et al., 2014). Long-term area change through time due to island ontogeny is not the only factor to influence island diversity and to vary throughout the island’s lifespan. At shorter-time scales, sea-level fluctuations also affect island area, as well as potentially altering island connectivity within an archipelago or with the mainland, by varying the distance between landmasses, for example through the formation of temporary land-bridges (Ali and Aitchison, 2014; Fernández-Palacios et al., 2016; Hammoud et al., 2021). Dynamic island connectivity has left signatures on contemporary total and endemic island species diversity (Weigelt et al., 2016; Norder et al., 2019). However, over macroevolutioanry timescales (*>* 1 million years), it is possible that rapid fluctuations may not leave a pronounced signature on phylogenetic and species richness data.

Most generalities of island biogeography derive from oceanic islands (Whittaker et al., 2008), limiting our understanding of continental islands, which, in fact, make up the vast majority of islands (Meiri, 2017). Oceanic islands initially completely lack terrestrial species, with diversity increasing through colonisation of mainland species. They tend to have a high proportion of endemics, given that many species undergo cladogenesis or anagenesis facilitated by island isolation (Valente et al., 2020). Cladogenesis on oceanic islands of sufficient area has led to spectacular radiations generally unrivalled in mainland ecosystems (Losos and Ricklefs, 2009). Continental island biota have distinct evolutionary characteristics. Often, a proportion of their species have vicariant origins, deriving from the mainland pool upon island formation. They generally have a lower proportion of endemics than oceanic islands (Ali, 2018). However, even within continental islands endemicity patterns vary strongly, with large old continental fragments such as Madagascar tending to harbour high levels of unique species, whereas recently formed land-bridge islands exhibit low endemism (Ali and Vences, 2019). Continental island systems, particularly those of small area, typically exhibit a decline in diversity right after their formation, a process termed ‘relaxation’, as the island is initially supersaturated (i.e. diversity exceeds the island’s area-dependent carrying capacity) (Diamond, 1972; Wilcox, 1978; Tilman et al., 1994; Kuussaari et al., 2009).

Testing the effect of relevant island characteristics, such as area, on island diversity through time is crucial to our understanding of island biogeography. Previous studies on the impact of ontogeny and fragmentation on islands have used correlational approaches (e.g. diversity and endemism data under space-for-time assumptions and species-area curves, He and Hubbell (2011); Damgaard (2019)). Simulation models that explicitly account for island ontogeny (Valente et al., 2014; Borregaard et al., 2016) or continental islands (Rosindell and Harmon, 2013) have provided insight into diversity dynamics but there is no likelihood-based inference framework yet for estimating parameters (colonisation, speciation and extinction) based on empirical data (Leidinger and Cabral, 2017). DAISIE (Dynamic Assembly of Island biota through Speciation, Immigration and Extinction) is a phylogenetic model of island biogeography developed to study oceanic island communities by inferring macroevolutionary processes. The use of phylogenetic information on colonisation times and speciation events allows accurate parameter estimation (Valente et al., 2018), but one may question to what extent these estimates are robust when the underlying process violates the model assumptions. For example, DAISIE has been applied to oceanic archipelagos assuming a constant archipelago area, arguing that submerging islands are replaced by emerging ones (Valente et al., 2015), but to date, the robustness of DAISIE under violations of its assumptions has not been tested.

Robustness analyses test the performance of a model when its assumptions are violated. These analyses provide insight into the generality of models by identifying when increased model complexity is unnecessary by testing model performance given departures from the model’s assumptions (Weisberg, 2006; Grimm and Berger, 2016). In model comparison there is no true model that perfectly represents reality (Burnham and Anderson, 2004). Therefore, finding simple models to explain complex dynamics can allow for inference in the presence of small sample sizes and reduce the risk of overfitting. In statistical modelling, it is possible to obtain reliable results when violating the assumptions of a model (e.g. t-test, Boneau (1960)). Testing inference robustness with simulations allows understanding of the type and magnitude of the violation, as long as the simulations are themselves without bias or applied in an unbiased fashion (Huelsenbeck, 1995; Hartmann et al., 2010). Robustness analysis is methodologically similar to testing model adequacy, which investigates the absolute performance of a model (Bollback, 2002; Pennell et al., 2015).

Here, we test the robustness of inference with a single model – DAISIE – to several geodynamic scenarios in which empirical data may differ from the model’s assumptions. We build upon the framework of Valente et al. (2014) to include ontogeny in the DAISIE simulations, and we simulate sea-level change, different types of island formation events (oceanic and continental), and transient land-bridge connection to the mainland. We then apply the maximum likelihood estimation (MLE) of DAISIE across this spectrum of geodynamic scenarios. Thus, we investigate to what extent a simple model of island biogeography can produce reliable results when ignoring geodynamics. If the model is found not to be robust, new inference methods that explicitly address geodynamics should be developed.

## Material and methods

### Oceanic area-dependent simulation model

The DAISIE framework consists of simulation models and inference models. The standard DAISIE simulation and inference model assumes that island communities assemble via cladogenesis (*λ^c^*), extinction (*µ*), anagenesis (*λ^a^*), and colonisation (*γ*), from a mainland species pool (of size *M*) (Valente et al., 2015). Cladogenesis and colonisation are the only rates assumed to be affected by a niche-filling process on the island, and can thus be modelled as diversity-dependent given a carrying capacity (*K’*) (Etienne et al., 2012; Valente et al., 2015) or as diversity-independent. Here we model diversity-dependence between species of the same clade (i.e. species stemming from the same mainland ancestor), but not between species of different clades.

In our oceanic area-dependent extension of the DAISIE simulation model, we model oceanic island area as a beta function of time, which allows various ontogeny scenarios (Valente et al., 2014), through four parameters: maximum island area, current island area, total island age, and the point in the island history where the maximum area is achieved. We model the relationship of cladogenesis and extinction rate with area as a power law (Valente et al., 2020). The parameters that define the power law of cladogenesis and extinction with area are denoted by *d* and *-x* respectively. Anagenesis rate is assumed to be independent of island area (Valente et al., 2020). Colonisation rate is dependent on area through the logistic term. Total rates of cladogenesis, extinction, colonisation and anagenesis at time *t* are calculated as follows using a time-dependent area, *A(t)*, given the number of species *N* at time *t*:

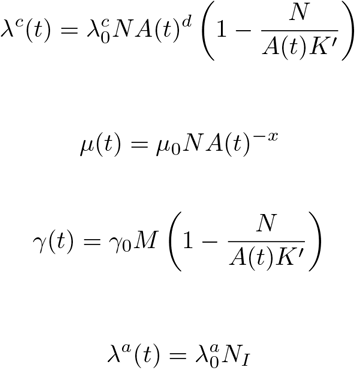

where *N_I_* is the number of immigrant (non-endemic) species on the island and 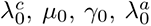 are the intrinsic rates of cladogenesis, extinction, colonisation and anagenesis, respectively, for unit area. The rates reduce to those of Valente et al. (2015), that is, the original DAISIE application, without area, when *d* = 0 and *A(t)* = 1 for cladogenesis, *x* = 0 for extinction, and *A(t)* = 1 for colonisation. *A(t)* is not needed if area is constant. We note that the functions that include a carrying capacity term (*K’*) reduce to diversity-independent functions when *K’* = *∞*.

Sea-level records show a semi-regular oscillation, so we model change in island area due to sea-level fluctuations as a sine wave (Miller, 2005; Bintanja et al., 2005). The sea-level model uses the same area-dependent evolutionary rates as above for island ontogeny. The sine wave is parameterised by amplitude and frequency. The effect of sea-level change on island area depends on the shape of the island. Here we assume that the island is a cone, where we set the slope. The area is calculated as the base of the cone at sea level, in accordance with other satellite island area measurements (Lim and Marshall, 2017; Valente et al., 2020), but a more realistic cone surface area can be used and is proportional to the base area (see Supplementary Methods).

### Continental island and land-bridge models

We extended the DAISIE simulation model further to model continental islands formed by separating from the mainland and remaining in isolation (“continental island” model) and continental islands that have transient connectivity as a result of the emergence and submergence of landbridges (“land-bridge” model).

The continental island simulation model uses the same core set of parameters as the original DAISIE simulation – *λ^c^*, *λ^a^*, *µ*, *γ*, *M*, and *K’* – with the addition of two sampling parameters: *x_s_* and *x_n_*. The number of species on the island upon its formation is determined by a sampling probability (*x_s_*), so the constant-rate oceanic simulation is a special case when *x_s_* = 0. As continental islands may have endemic species upon formation, e.g. range restricted species, the endemicity of each species sampled at island formation is determined by the non-endemic sampling probability (*x_n_*). Island formation is assumed to occur instantaneously. For this study, we assume constant-area for continental islands to allow us to focus on the effect of the change in connectivity of the island without the other confounding area-related effects. For the effects of the latter, we refer to the oceanic island scenarios.

The land-bridge simulation model is fundamentally a birth-death-shift model (Rabosky, 2006; Stadler, 2011; Höohna et al., 2016), in which parameter rates can shift instantaneously to another set of time-constant rates in a piecewise manner (in our case *λ^c^*, *λ^a^*, *µ*, *γ*, *K’*). In the simulation, land-bridge formation is not determined by sea-level fluctuations, but is instead modelled phenomenologically. The island has two rates: one for periods when the island is separated from the mainland and another for periods when the island is connected by a land-bridge to the mainland. For the land-bridge model, we assume area is constant in both connected and unconnected states. Rates can be diversity-dependent or diversity-independent. The shift between the two states can occur any number of times, with a single shift model (Valente et al., 2019; Hauffe et al., 2020) being one scenario. DAISIE can already make inferences using a single shift-rate model to empirical data (Valente et al., 2019; Hauffe et al., 2020); however, given the frequency of land-bridges over macroevolutionary time frames (e.g. multiple glacial cycles), testing robustness to multiple shifts is still warranted.

### Simulating phylogenetic data with geodynamics

We simulated island biogeographic processes using a constant-rate or time-dependent Doob-Gillespie algorithm (Gillespie, 1976; Allen and Dytham, 2009) implemented in the R package DAISIE (Etienne et al., 2021). The evolutionary history of each species on the island is recorded and phylogenetic trees for island communities are produced. Henceforth, the original DAISIE simulation (Valente et al., 2015) is termed *constant-area oceanic simulation* (excludes ontogeny, sea-level fluctuations and continental scenarios), whereas the new simulations introduced in this paper are termed *geodynamic simulations* (island ontogeny, sea-level changes, continental and continental land-bridge). Given the total number of possible combinations of geodynamic simulation scenarios, a complete exploration of the parameter space for each simulation model would be computationally unfeasible. Therefore, we restricted our parameter space to realistic parameter values inspired by empirical studies (supplementary tables S1-S9).

The parameter space for the area-dependent models encompasses a realistic range of cladogenesis, extinction, colonisation and anagenesis; while the carrying capacity ranges across realistic finite values (diversity-dependent) and an infinite value (diversity-independent) (Valente et al., 2020) (supplementary tables S1-S6). For the island ontogeny model, we used parameters for area curves from the geological literature to model a young and an older island. These parameters were inspired by Maui Nui and Kaua’i from the oceanic archipelago Hawaii (Lim and Marshall, 2017). These two islands provide contrasting ontogenies, one – Maui Nui, the “young island” – with a shorter geological time span (2.55 My), and sharp increase to a large maximum area, and the other – Kaua’i, the “old island” – with a longer geological time span (6.15 My) and shallower increase to a smaller maximum area (Fig. 1). We assume that ontogeny describes a single island (which is not the case for Maui Nui as it currently includes four main islands, which were previously a single island). We simulated from the islands’ origin to the present. We used sea-level data from the last one million years to calibrate the area curve for sea-level (supplementary tables S3-S6) (Bintanja et al., 2005). For each sea-level scenario, we simulated two different island gradients: 80*^◦^*and 85*^◦^*, and thus have two different magnitudes of area change through time for a single sea-level fluctuation through time. When only sea-level dynamics are present (no ontogeny), we assumed sea-level fluctuates around the current island area for the young and older island.

**Figure 1:**
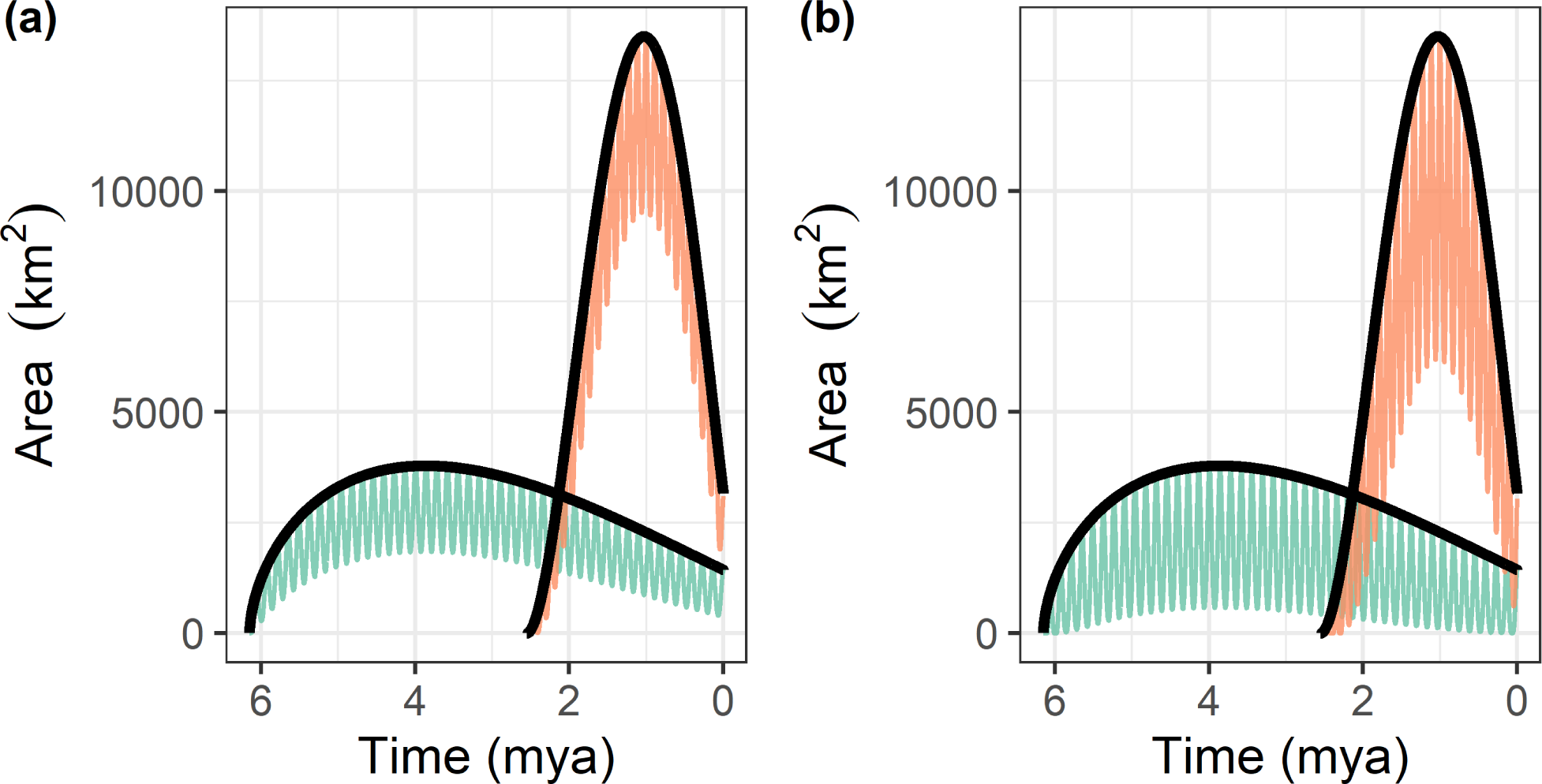
Variation in island area through time simulated in oceanic ontogeny and oceanic ontogeny sea-level scenarios. The black lines correspond to area variation through time from oceanic ontogeny simulations, calibrated to the island age, maximum age and time of peak area for the young and old islands. The orange and green lines correspond to area variation through time from oceanic ontogeny and sea-level simulations for the young (orange) and old (green) islands. In (a), the variation in island area in the oceanic ontogeny sea-level scenario is shown for a steep (85*^◦^*) island and following the same geological calibration as above. Because the modelled islands are steep, sea-level fluctuations have a lower impact on island area. In (b), the same scenarios are presented, but here shallow islands (80*^◦^*) are modelled, resulting in a greater influence of sea level on island area. mya – million years ago.

We tested the continental island model using two sampling probabilities (*x_s_*) and non-endemic sampling probabilities (*x_n_*) (supplementary tables S7-S9). For the land-bridge model we simulated the emergence and submergence of one land-bridge. Anagenesis is fixed to 0 when in the land-bridge state, as this mode of speciation is meaningless given the land-bridge connection. To test the effect of land-bridge properties on DAISIE we varied the colonisation rate when the land-bridge is present (land-bridge colonisation multiplier, i.e., the factor by which the colonisation rate increases when the land-bridge is present) (supplementary tables S8-S9). Land-bridge rates of cladogenesis and extinction are assumed to be half the island rates due to the increased gene flow and rescue effect (Brown and Kodric-Brown, 1977; Rosindell and Phillimore, 2011). We also simulated a constant-area young and an old island for the continental island and land-bridge models. Additionally, for continental islands we also simulated an ancient island, to mimic ancient continental fragments such as Madagascar or New Zealand. In this case we used a much older island age of 50 My in order to test whether the signature of species initially present when the island is formed decays through time and the inference error decreases when DAISIE is applied to the data.

### Robustness pipeline

We developed a novel pipeline to analyse the robustness of a phylogenetic model to any simulation (Fig. 2). The pipeline quantifies the error made when inferring processes from data simulated with complexities not included in the inference model.

**Figure 2:**
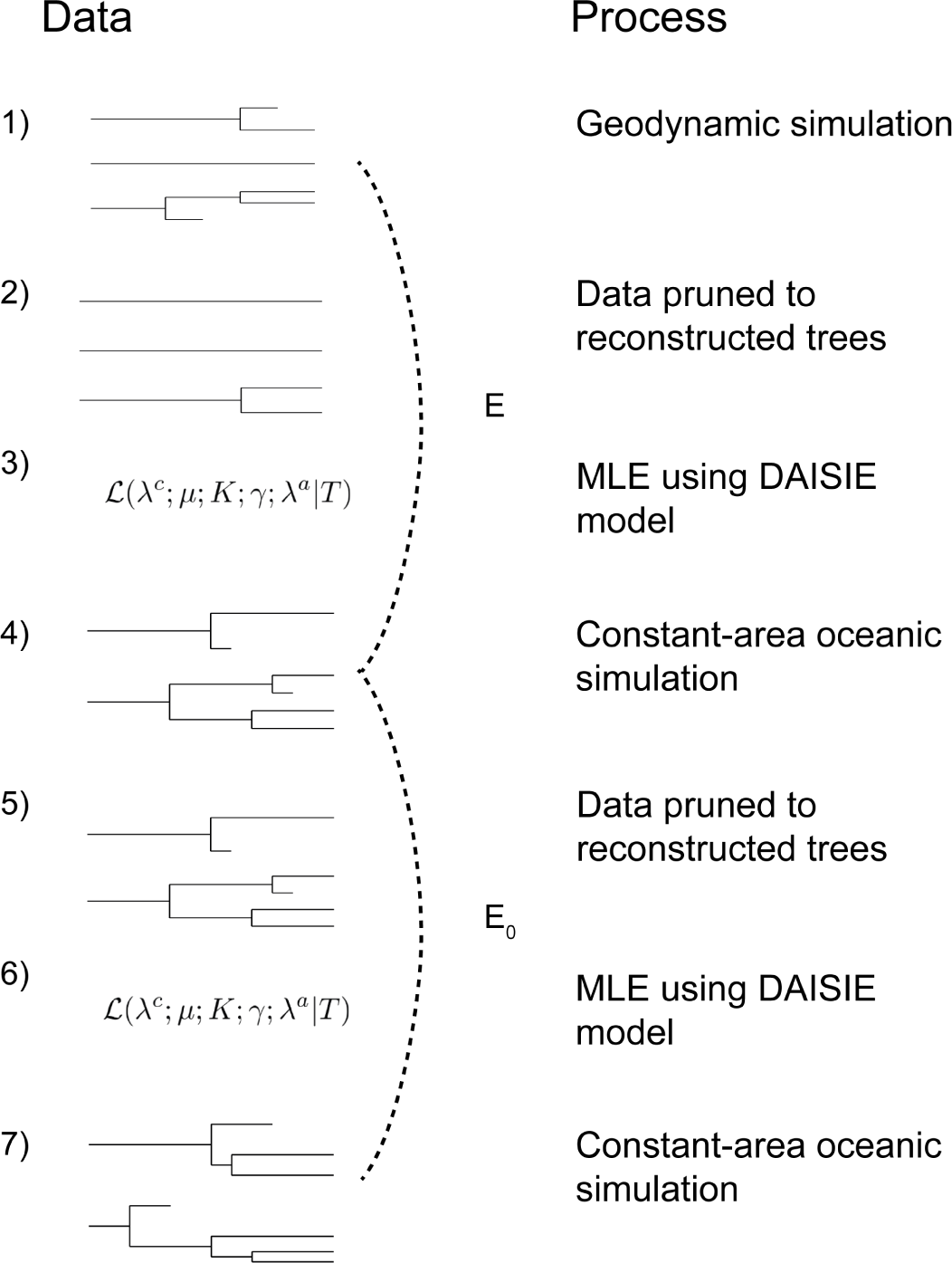
Schematic representation of the robustness pipeline. (1) Island phylogenetic data is produced by a geodynamic simulation. (2) Reconstructed phylogenetic data is produced by pruning (removing) all extinct lineages. (3) DAISIE maximum likelihood estimation (MLE) of parameters, which assumes no geodynamics, is applied to data from step 2. (4) A constant-area oceanic simulation (i.e. the original DAISIE method) is run with the parameters estimates from step 3 as generating parameters. (5) Phylogenies are pruned to reconstructed trees. (6) The DAISIE MLE is applied to data from step 5. (7) A constant-area oceanic simulation is run with parameter estimates from step 6 as generating parameters. The inference error made on data from a geodynamic simulation (E, step 3) is compared with the baseline error (E_0_, step 6). The error E is calculated by comparing the geodynamic simulations with the first set of constant-area oceanic simulations for five metrics (dashed line). The baseline error E_0_ is obtained by comparing two oceanic simulations using the same five metrics (dashed line). The error made when inferring from geodynamic data but assuming the constant-area oceanic model in inference is the proportion of errors (E) that exceed the 95th percentile of the baseline errors (E_0_).

The geodynamic simulation models were iterated for 1,000 replicates for every parameter set to account for stochasticity. We conditioned on five colonising lineages surviving on the island to the end of the simulation. This ensured that the MLE model uses data with a sufficient sample size to reliably estimate parameters, while also being representative of previous empirical data applications of the MLE model (Valente et al., 2018). Parameter estimation was then carried out on each replicate using the oceanic DAISIE MLE model with diversity-dependence (Valente et al., 2015). The likelihood is conditioned on five colonising species surviving to the present to be consistent with the conditioning implemented in the simulations. We then used the parameters inferred from the MLE to simulate under the constant-area oceanic simulation model (which is the same as the DAISIE inference model), for a single replicate per geodynamic replicate with both simulations having an equal length of time (Fig. 2).

### Inference error

In order to determine the robustness of the DAISIE inference model to data simulated under geodynamics, we calculated the inference error on geodynamic data (E) and compared that to the estimation error inherent in the DAISIE maximum likelihood (baseline error, E_0_). The former is the error when violating DAISIE’s assumptions, and the latter is the intrinsic inference error when the DAISIE assumptions are met (i.e. the simulation model is identical to the inference model).

We quantified the error using the ΔnLTT statistic (Janzen et al., 2015) and two island diversity metrics, the number of species and number of colonisations at the present. The ΔnLTT statistic provides a non-parametric approach that quantifies the divergence between two islands in the number of species-through-time. It can be applied to both reconstructed and full phylogenies, allowing for quantifying differences even when clades are in decline, which is rarely detected in models fitted to reconstructed trees (Burin et al., 2019). The ΔnLTT statistic is defined as the difference in normalized lineage-through-time (nLTT) between the geodynamic simulations (Fig. 2 step 1) and the constant-area oceanic simulations generated with the MLE estimates inferred from the geodynamic simulations (Fig. 2 step 4). We applied the ΔnLTT statistic to the total species-through-time (ΔSTT), endemic-species-through-time (ΔESTT) and non-endemic-species-through-time (ΔNESTT) for each data set. All error analyses used the DAISIErobustness (Lambert et al., 2021) and the DAISIE (Etienne et al., 2021) R packages using R version 4.1.0 (R Core Team, 2021).

From the divergence between the error and the baseline error we calculated a test statistic which we denote by ED_95_. This is defined as the number of replicates in which the error is greater than the 95th percentile of the baseline error (R_95_) plus one, divided by the number of replicates (*N*) plus one: (R_95_ + 1)/(*N* + 1). When the error distribution largely overlaps with the baseline error distribution (Fig. 3a) the ED_95_ statistic is low (i.e. for complete overlap ED_95_ = 0.05), whereas in cases when the error distribution is mostly divergent from the baseline error distribution (Fig. 3b) the ED_95_ statistic is higher (i.e. for complete divergence ED_95_ = 1).

**Figure 3:**
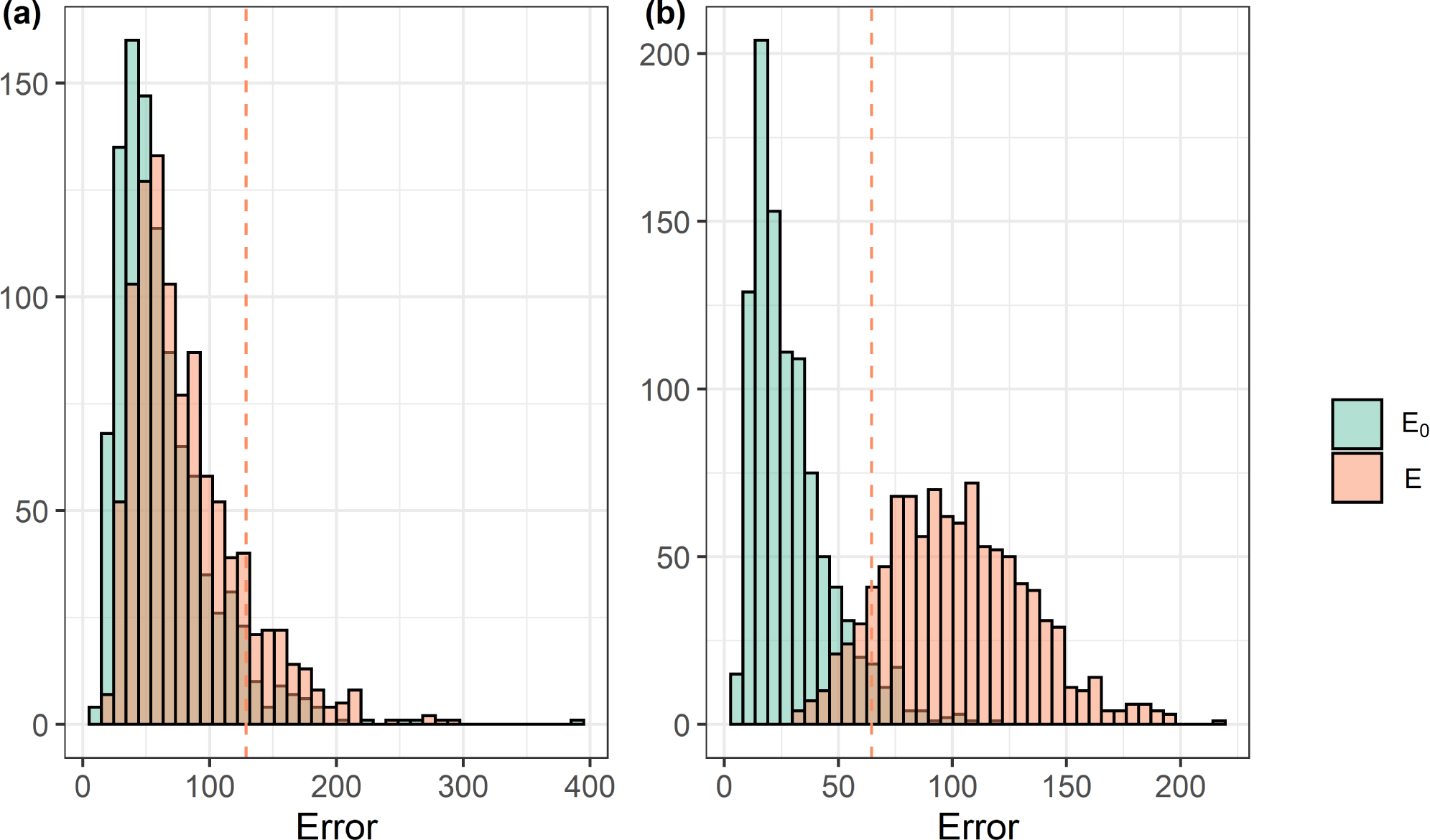
Hypothetical examples of histograms of error (E) and baseline error (E_0_) distributions for two parameter sets. The dashed pink line corresponds to the 95th percentile of the E_0_ distribution. In (a) the E and E_0_ distributions are largely overlapping and a large proportion of the E_0_ distribution falls to the left of the dashed pink line (low ED_95_ value). By contrast, in (b), the majority of the error distribution falls to the right of the dashed pink line (high ED_95_ value).

The robustness pipeline (Fig. 2) was run until 1,000 replicates were complete for each parameter set. In some cases, 1,000 replicates took an exceedingly large amount of time. As a result, we limited the computational run time to ten days. Within ten days most parameter sets finished (oceanic ontogeny: 371 out of 384 (97%); oceanic sea-level: 750 out of 768 (98%); oceanic ontogeny sea-level: 758 out of 768 (99%); continental island: 230 out of 576 (40%); land-bridge: 148 out of 256 (58%). There was no correlation between computation time and robustness for the parameter sets that finished within ten days and thus we assume there is no bias in the results presented below (Fig. S1).

## Results

### Geodynamic oceanic islands

Oceanic islands with ontogeny, sea-level fluctuations, or both cause minimal error when reconstructing total species, endemic species, and non-endemic species through time based on parameters inferred with the non-geodynamic DAISIE inference model, (Fig. 4 & 5, Fig. S2 & S3). The error is controlled by the strength of the effect of island area on cladogenesis and extinction rates (Fig. 5 & Fig. S2 & S3). The number of species and island colonising clades at present are both remarkably well predicted (Fig. 4g & 4i). In other words, DAISIE inference performs well when applied to oceanic islands with geodynamic area. Hyperparameters (*x* and *d*) controlling the effect size of area change determined the error made on ΔSTT and ΔESTT for oceanic ontogeny (without sea-level) with high values of *d* (strong effect of area on cladogenesis) causing the largest error, especially for the old island (Fig. 5a, Fig. S2a & S2c). For sea-level change on its own, the error in ΔSTT and ΔESTT was around the null expectation (0.05) for all hyperparameters for both island ages (Fig. 5b & Fig. S2b). In the case of ontogeny and sea-level operating together, inference with the non-geodynamic DAISIE model showed higher error in ΔSTT compared to ΔESTT and ΔNESTT (Fig. 5). When reconstructing non-endemics through time, oceanic island ontogeny and sea-level fluctuations, separately, cause minimal error on DAISIE inference, which is not strongly affected when altering the effect of area on cladogenesis (*d*) or extinction (*x*) (Fig. S3a & S3b). Ontogeny and sea-level operating together produce somewhat higher error in ΔNESTT (Fig. S3c). The ΔSTT, ΔESTT, and ΔNSTT have equal error at both island gradients for sea-level (Fig. S4) and ontogeny plus sea-level (Fig. S5).

**Figure 4:**
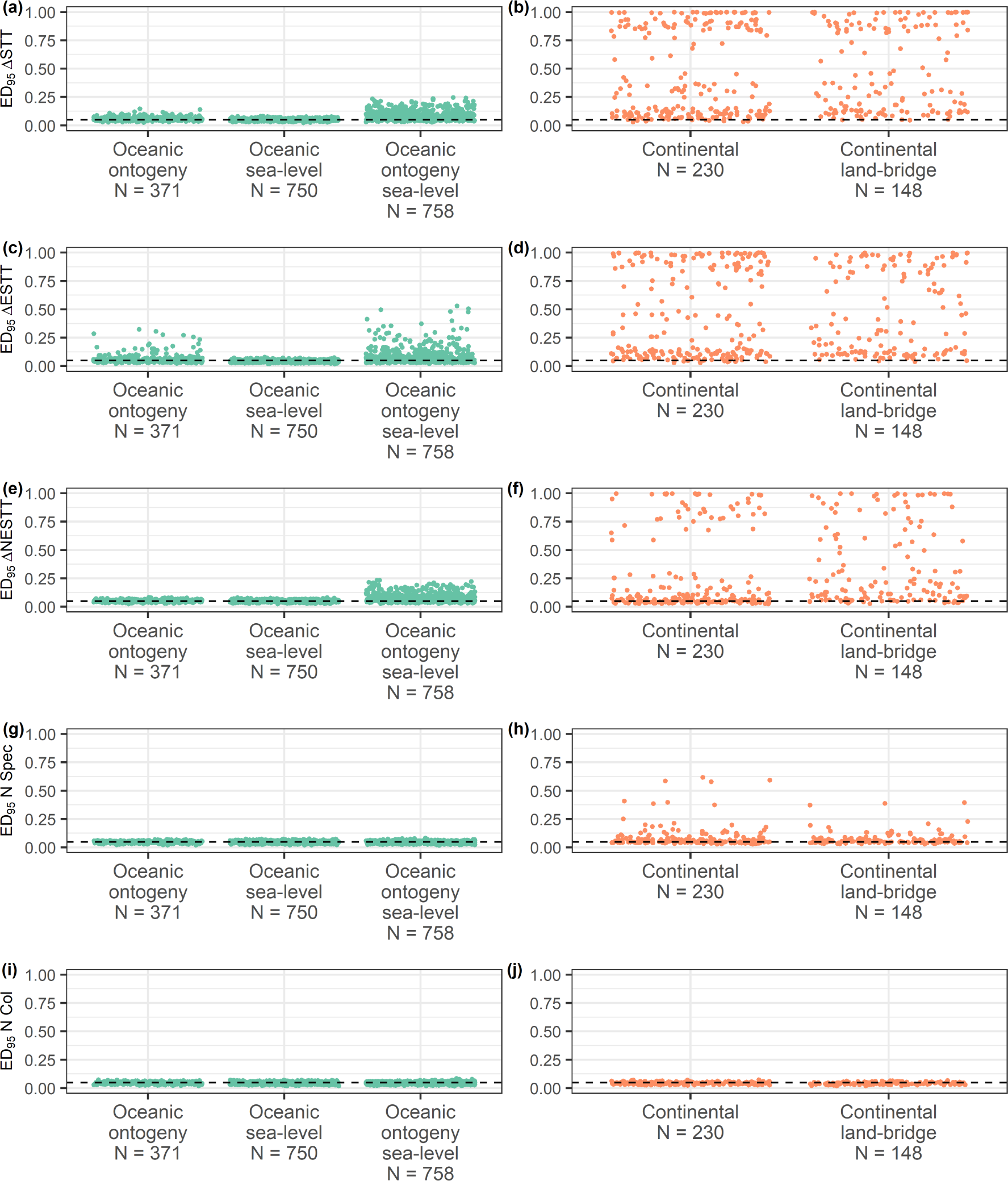
Strip charts showing the distribution of the ED_95_ statistics calculated for the ΔSTT (a-b), ΔESTT (c-d), ΔNESTT (e-f), number of species at the present (N Spec) (g-h), and number of colonists at the present (N Col) (i-j) for each geodynamic scenario. The scenarios are: oceanic ontogeny, oceanic sea-level, and oceanic ontogeny sea-level (green), as well as two continental scenarios: continental island and continental land-bridge (orange). Each point represents the ED_95_ for a single parameter setting. Dashed line at 0.05 is the null expectation of the ED_95_ error. N shows the sample size for each strip on the *x*-axis.

**Figure 5:**
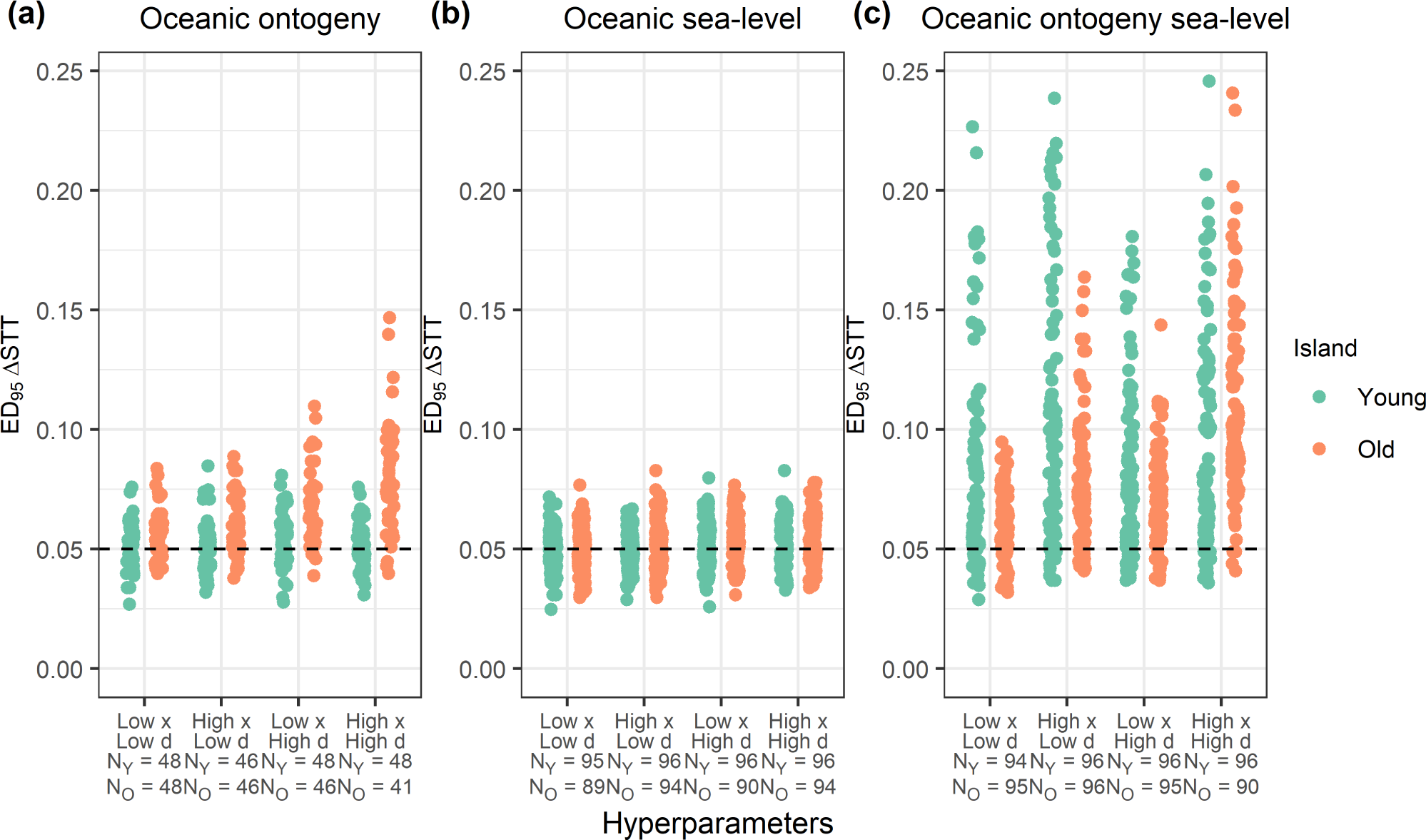
Strip charts showing the distributions of the ED_95_ statistic for ΔSTT for each combination of hyperparameters (*d* and *x* controlling the effect of area on the rates of cladogenesis and extinction respectively) for the dynamic area oceanic island scenarios. Each point represents the ED_95_ for a single parameter set with the specified hyperparameters on the *x* -axis. All plots have a dashed line at 0.05 which is the null expectation of the ED_95_. (a) ΔSTT ED_95_ statistic for oceanic ontogeny. (b) ΔSTT ED_95_ statistic for oceanic sea-level. (c) the ΔSTT ED_95_ statistic for oceanic ontogeny and sea-level. The sample size for each strip is shown for young islands (N_Y_) and old islands (N_O_). Note that the *y*-axis scale is a subset of that in Figure 4, to allow better visualization of small ED_95_ values.

The number of species and number of colonists at the present for all oceanic area-dependent scenarios showed negligible error, with all results clustering around the expected baseline error from the null prediction (i.e. no additional error, ED_95_ *≈* 0.05) (Fig. 4d & 4e). Changes in hyperparameter values (*d* and *x*) and island gradient (for the scenarios that include sea-level) did not considerably alter the error for the number of species and colonists (Fig. S6-S9).

### Continental islands with and without land-bridges

In the scenarios in which the island inherits species upon formation (continental island) and has intermittent connections to the mainland (land-bridge) the DAISIE ML inference based on a constant-area oceanic model is often unable to accurately reconstruct diversity through time, and frequently shows vastly greater error than oceanic area-dependent scenarios (Fig. 4). The effect of having species initially present on the island causes error in the estimation and reconstruction of the island diversity through time, and this error is proportional to the initial number of species on the island (*x_s_*) (Fig. 6). The impact of having species initially on the island is mediated by the island’s lifespan for reconstructing species, endemics and non-endemics through time (Fig. 6). For young islands, even a few initial species cause error in species and endemics through time (Fig. 6a & 6b). The error in reconstructing non-endemic species is determined by both *x_s_* and *x_n_*; when both are high a large amount of error is produced, but when either are low the error is also relatively small (Fig. 6c). In all continental scenarios (i.e. all combinations of *x_s_* and *x_n_* excluding land-bridge scenarios) the ancient islands (simulated for 50 Myr) are relatively robust, indicating the signature of continental islands is eroded as the island remains isolated for longer periods of time (Fig. 6).

**Figure 6:**
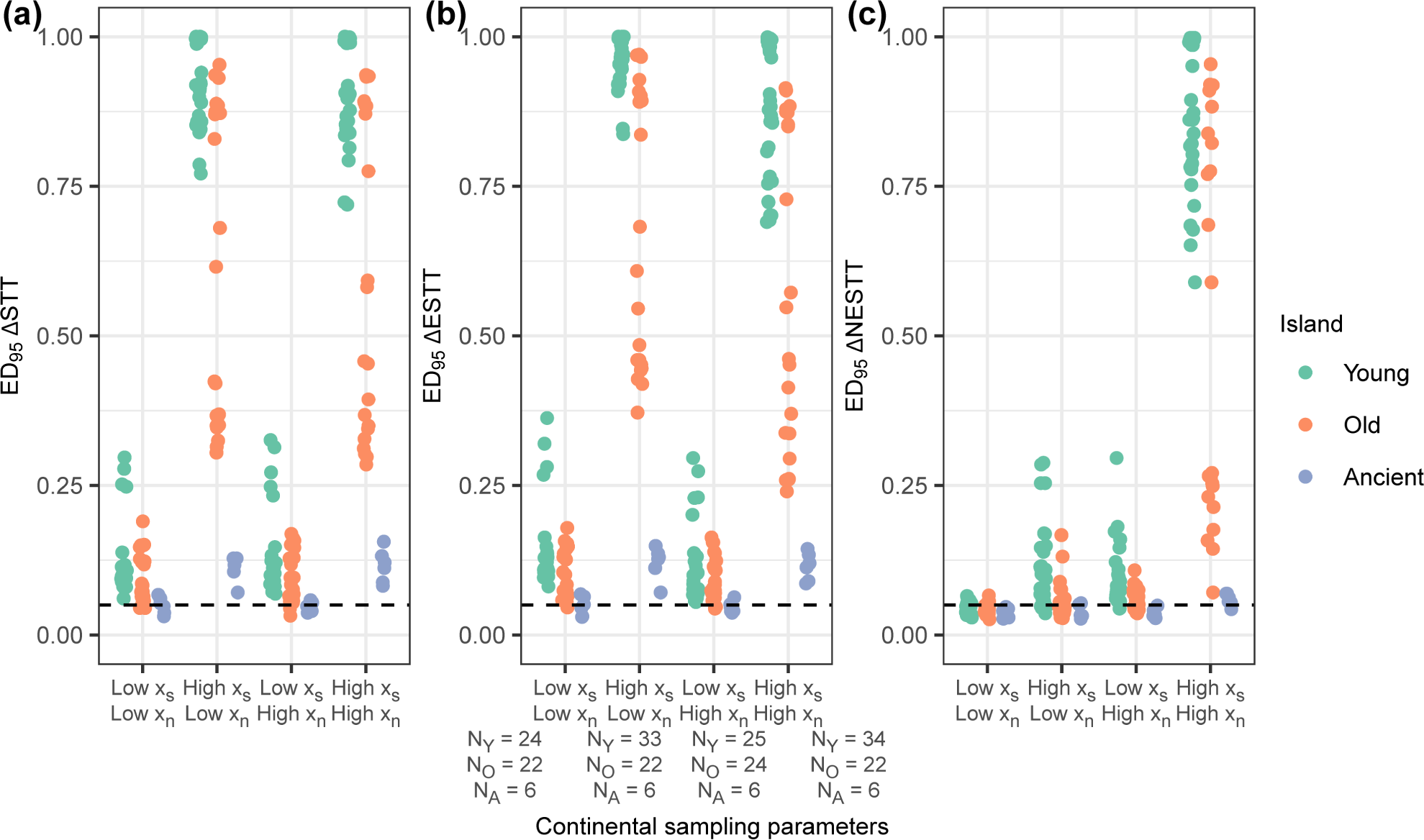
Strip charts showing the distributions of the ED_95_ statistic across the combinations of continental sampling parameters (*x_s_* and *x_n_*), for each island age. Each point represents the ED_95_ for a single parameter set with continental sampling parameters on the *x* -axis. All plots have a dashed line at 0.05, which is the null expectation of the ED_95_. Metrics plotted are: (a) ΔSTT ED_95_ statistic, (b) ΔESTT ED_95_ statistic, and (c) ΔNESTT ED_95_ statistic. The sample size for each strip is shown for young islands (N_Y_), old islands (N_O_), and ancient islands (N_A_), sample size for (a), (b) and (c) are the same.

The magnitude of the error made in the continental land-bridge model is roughly equal to continental islands without land-bridges across species, endemics and non-endemics through time (Fig 4a-4c, Fig. 6 & Fig. S10-S12). The influence of different immigration rates when the land-bridge is present does not drastically alter the error (Fig. 7). The results for endemic species-through-time (Fig. 7b) are similar to species-through-time (Fig. 7a). For the young island simulations, when the number of species initially on the island is low, there is little error in the number of endemics through time. Alternatively, when the initial number of species is high, there is a large amount of error in endemics-through-time. When the number of endemics initially on the island is high (*x_n_* is low), the error in endemic species-through time is large. The error from continental island land-bridge was similar across ΔSTT, ΔESTT and ΔNESTT (Fig. S10).

**Figure 7:**
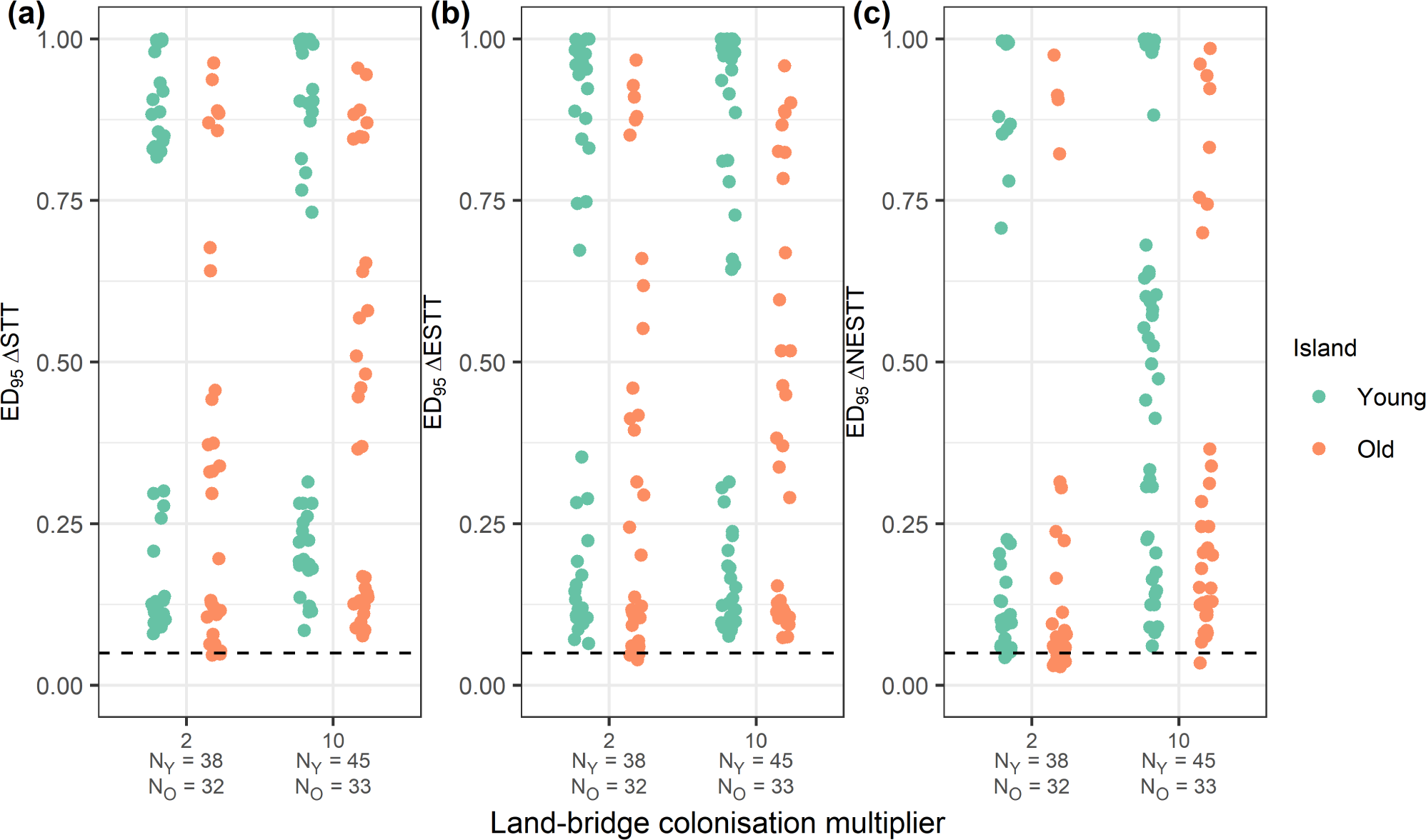
Strip charts showing the influence of different land-bridge colonisation multipliers (i.e., the factor by which the colonisation rate increases when the land-bridge is present). Each point represents the ED_95_ for a single parameter set. All plots have a dashed line at 0.05 which is the null expectation of the ED_95_. Metrics plotted are: (a) ΔSTT ED_95_ statistic, (b) ΔESTT ED_95_ statistic, and (c) ΔNESTT ED_95_ statistic. The sample size for each strip is shown for young islands (N_Y_) and old islands (N_O_).

For continental islands, when species are initially on the island, the present-day species diversity and number of colonists is accurately reconstructed in most cases, with or without the presence of land-bridges (Fig. 4d-4e, S11 & S12). Continental young islands show that under certain continental island parameter settings the DAISIE model cannot reconstruct diversity to present-day levels very well (Fig. S12a). For continental land-bridge scenarios the exception from negligible error is the number of species for the old island (Fig. S10a).

## Discussion

Our analysis shows that a phylogenetic model of island biogeography that does not account for dynamic island area can accurately reconstruct several metrics of biodiversity through time for islands that form *de novo*, i.e. oceanic islands. However, the same model is usually not robust to the presence of species at island formation, a scenario likely in islands that break away from the continent or are connected to the continent by lowlands that may be flooded during interglacial periods. The age of separation of the island from the mainland is a key indicator for robustness of DAISIE to continental islands – the longer the island has been separated, the higher the robustness of DAISIE.

Traditionally, models that estimate macroevolutionary dynamics from reconstructed phylogenies (i.e. only extant species) can only analyse a single phylogeny, excluding species-poor clades due to insufficient data. By contrast, DAISIE is a phylogenetic model that estimates the macroevolutionary dynamics for an entire community of island species (composed of multiple independent “phylogenies”, many of which with only a single species). This facilitates the investigation of questions that were previously difficult to address on evolutionary time scales in the field of island biogeography, for example whether equilibrium or non-equilibrium dynamics operate on an island (Valente et al., 2015). The precision of the model has been validated when the inference process is matched by the generating process (Valente et al., 2015, 2017a, 2018), and in this study the model has been shown to accurately infer and reconstruct island diversity patterns when influenced by area.

The lack of robustness of DAISIE to continental islands only applies to the parameter space that was tested within this study, with a conservative estimate of the probability of vicariant species on the island (0.01 and 0.05). Continental islands in nature may exceed this, but there is little information on the number of species that were initially present on a continental fragment or land-bridge island after isolation, as well as what proportion of those species were endemic.

### Robustness to area changes on oceanic islands

The idea of biodiversity on islands being in disequilibrium has been theorised (Whittaker et al., 2008; Fernández-Palacios et al., 2016) and the impact of sea-level change and ontogeny on island diversity and endemism has previously been shown (Lim and Marshall, 2017; Norder et al., 2019). Recent attempts at understanding the macroevolutionary dynamics from phylogenetic data have not considered these geological and sea-level changes when making claims about island community assemblage and diversification (Valente et al., 2015, 2020). Here we show that although in reality island communities may be governed by geological and eustatic characters, a simple phylogenetic model of island biogeography – DAISIE – can reconstruct diversity dynamics (number of species at present and how they vary through time, as well as the numbers of colonists at present) with little error. The minimal error made by DAISIE when inferring parameters from data produced under realistic island ontogeny of two Hawaiian Islands supports the reliability of previous results using DAISIE. As an example, the Galápagos and Macaronesian archipelagos have complex geographic histories, experiencing rapid island formation from a mantle plume, followed by slow subsidence, and both systems have undergone area changes due to sea-level fluctuations (Fernández-Palacios et al., 2011; Ali and Aitchison, 2014; Geist et al., 2014; Rijsdijk et al., 2014). DAISIE has been applied to the avifauna of the Galápagos and Macaronesia assuming the current archipelago geography back to the system’s origin. The island ontogeny with sea-level changes within this study resembles the histories of such archipelagos, and by extension indicates the reliability of DAISIE’s previous findings on dynamic oceanic systems. The simulations to test geodynamics do not account for the fact that islands within an archipelago may have been connected and disconnected (Aguiĺee et al., 2021). However, under the assumption that the archipelago operates as a single island – an assumption most valid for volant species e.g. avifauna of the Galápagos (Valente et al., 2015) and Macaronesia (Valente et al., 2017b) – the change in area through time can mimic archipelago dynamics.

This study has shown that oceanic island biodiversity governed by time-dependent rates can be approximated with little error even without knowledge of time-dependent rates. The development of an inference model of island biogeography that includes changes in area and changes in isolation could be accomplished, with time-variable rates regularly used in phylogenetic models (Condamine et al., 2019). The problematic aspect is the risk of overfitting a complex model to information-poor data sets, with oceanic islands typically hosting communities between 10 and 50 species for a given taxon of interest (e.g. birds, Valente et al. (2020)), even when they are typically diverse on the continent. The results we find here also suggest that area-dependent rates do not leave a significant signature on phylogenetic data, and as a result, a new inference model incorporating changes in area may not have adequate information to estimate such time-variable dynamics, and may thus erroneously predict such dynamics.

The robustness of DAISIE shown in this study does not extend to making predictions about the (distant) future of island biodiversity. If a model does not accurately incorporate geodynamics, using estimated parameters to make predictions about island biodiversity into the future will be biased (Valente et al., 2017a). For example, the constant-area oceanic DAISIE will predict species can exist forever, even on oceanic islands which will eventually subside. The robustness found within this study for oceanic islands may be due to island diversity not declining as fast as expected in nature. The relationship between extinction and island area is inspired by the species-area relationship historically found to be a power law (Dengler, 2009). Valente et al. (2020) showed that the relationship between island area and extinction was better described by a power law compared to a sigmoidal relationship, but extensions of the power law have not been explored, with other formulations resulting in fewer species at smaller areas (Plotkin et al., 2000).

### Vicariant and land-bridge species cause poor performance

The largest error when reconstructing several aspects of diversity through the island’s history from DAISIE estimates occurs under continental scenarios. This error was determined by the number of species initially on the island and the time since the island’s formation, with the robustness of DAISIE increasing as the number of initial species decreased and the time of island separation from the mainland increased. The lower error for longer island history in the continental island case is likely a result of species turnover erasing the signature of vicariant species initially on the island. In these cases, the longer the island exists, the larger the proportion of species that come from colonisation events from the mainland and more of the initial biota go extinct, and the island becomes quasi-oceanic from a biodiversity perspective. Additionally, empirical evidence indicates that heightened extinction from relaxation on continental islands will erase much of the signal of vicariant species over evolutionary time scales, which would produce reconstructed phylogenies resembling oceanic islands (Diamond, 1972; Halley et al., 2016). A limitation of note here is that our simulation model does not produce characteristic periods of relaxation after island formation, as species from different clades on the island do not compete and colonisation plus cladogenesis balance extinction. A model in which all species on the island compete (Etienne et al. *in prep*) or without colonisation or cladogenesis (Halley and Iwasa, 2011) could produce more realistic biodiversity dynamics for after continental island formation.

DAISIE has been applied to the continental archipelagos of New Zealand (Valente et al., 2019) and the Greater Antilles (Valente et al., 2017a). New Zealand is estimated to have been isolated from other landmasses for at least 52 million years (Schellart et al., 2006), which is a much longer period of time than the age of most volcanic oceanic islands (Valente et al., 2020). Wallace (1880) stated that New Zealand’s biota is akin to that of an oceanic island, an idea that our results for ancient continental islands seem to support. The Greater Antilles is an archipelago consisting of multiple ancient continental fragments and may have been last connected to continental America by a land-bridge at the Eocene-Oligocene transition (35-33 million years ago, Iturralde-Vinent and Macphee (1999)). While there are terrestrial species of vicariant origin in the Greater Antilles (Brace et al., 2015), phylogenetic data for bats supports overwater dispersal for all extant and recently extinct taxa (Valente et al., 2017b). Therefore, our results suggest that previous DAISIE conclusions on ancient continental fragments are reliable. A model that takes into account species initially on the island is theoretically possible within the DAISIE framework, and can be explored for recent continental islands which show high error in this study.

The theory of species dispersal controlled by geological evolution, and specifically land-bridge formation and disappearance was laid out by Simpson (1940). Since then it has become clear that land-bridges facilitate dispersal and alterations in species ranges (Wilcox, 1978). In the land-bridge simulations studied here we also found that higher colonisation can cause shifts in the island community which cannot be accounted for by the simple inference model. In practice, DAISIE has not been applied to islands that have experienced recent land-bridge formation, because it was not designed for this. The error made on land-bridge islands exemplifies the need to apply models that account for pairwise shifts in community rates (Hauffe et al., 2020). The failure of DAISIE to robustly estimate rates of macroevolution on land-bridge islands shows clear evidence that oceanic and continental islands leave distinct macroevolutionary signatures in phylogenetic data and thus different models should be used to study their dynamics. Indeed, phylogeographic and paleo-island biogeography studies have revealed that land-bridge islands have unique colonisation, migration and speciation dynamics that often do not fit classic oceanic-island-centric island biogeography (e.g. Papadopoulou and Knowles (2015); Hammoud et al. (2021)). The land-bridge implementation with different rates of land-bridge colonisation used in this study is consistent with how different taxonomic groups may utilise land-bridges to colonise at different rates. For example, in the case of terrestrial organisms (e.g. amphibians and mammals), the difference in rates between periods of land-bridge emergence and submergence will likely exceed the difference found in birds, which may have a higher background rate of colonisation when the land-bridge is absent. The different land-bridge colonisation rates are also compatible with different modes of increased colonisation to an island for a period of time, for example the low colonisation multiplier can represent periods of increasing island colonisation via oceanic currents facilitating rafting (Ali and Huber, 2010), whereas the higher colonisation multiplier may represent a physical land-bridge aiding range movement onto islands.

### Robustness studies in phylogenetics

Testing the performance of a model of species diversification using reconstructed phylogenies, either by testing model adequacy or the type I error rate and power, has previously been explored (Davis et al., 2013; Pennell et al., 2015; Rabosky and Goldberg, 2015; Etienne et al., 2016). However, these studies assume the inference model is the same as the generating model used to simulate the data. Investigating the behaviour of an inference model when the generating model is different has not been frequently used in models fitted to phylogenetic data (but see Simonet et al. (2018)). On the other hand, the robustness of models of phylogenetic inference has been tested and is well established (Huelsenbeck, 1995; Bilderbeek et al., 2021). Interestingly, these studies have shown that under certain conditions a bias favours simple models over more complex alternatives (Yang, 1997; Bruno and Halpern, 1999). The robustness of the DAISIE inference model to violations in its assumptions highlights the justification for general approximations to describe complex phenomena with a high number of degrees of freedom. Biological systems of interest are highly complex, with heterogeneity between individuals scaling up to macroevolutionary dynamics. Therefore, as in other fields of science, approximations are warranted for a generalised understanding of a system, but should be justified through analysis of a model’s robustness to violations. The generality of the robustness pipeline developed here allows for other studies on the robustness of other models under complex generating processes. Alternative hypotheses for island changes are changing isolation, topographical complexity, or evolution of environmental heterogeneity as new islands undergo succession (Massol et al., 2017; Barajas-Barbosa et al., 2020).

## Acknowledgments

We would like to thank Shu Xie, Giovanni Laudanno, Richèel J.C. Bilderbeek, and Pratik Rajan Gupte for their input. We would like to thank the Center for Information Technology of the University of Groningen for their support and for providing access to the Peregrine high performance computing cluster. PSN was funded through a FCT PhD Studentship with reference SFRH/BD/129533/2017, co-funded by the Portuguese Ministéerio da Cîencia, Tecnologia e Ensino Superior and the European Social Fund. JWL was funded through a Study Abroad Studentship by the Leverhulme Trust and was also funded by a NWO VICI grant awarded to RSE. LV was funded by a NWO VIDI grant.

## Supplementary Material

### Supplementary Methods

#### Sea-level area changes calculations

Sine wave (*SW*) for sea-level fluctuations:

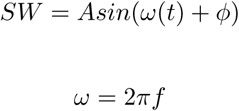

Surface area (*A*) of a cone:

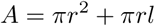

where *l* is the length of the hypotenuse of the cone, *h* is the height of the cone, and *r* is the radius of the cone.

For the surface area of the upper surface of the cone we drop *πr*^2^.

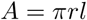

Express the length *l* in terms of *r* and replace *l* in the surface area equation:

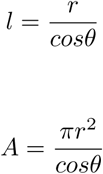

Calculate the radius *r* of the cone by rearranging:

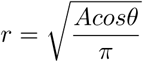

Calculate the height *h* from the radius *r* and the tangent of the angle:

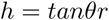

Expressing *r* and *l* in terms of *h* we can write area as:

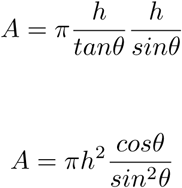

Given the current height of the cone (*h*_0_) and the change in the height given a change in sea-level (*h*) the new surface area of the cone is:

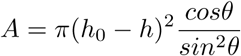

Area of the base of the cone instead to model the projection area (area as measured from an aerial view) commonly used in island biogeography.

Base of the cone:

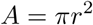

Re-write with *r* in terms of *h*:

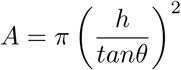

Given sea-level change (*h*):

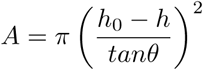

Therefore, the relationship between surface area of the cone and the base of the cone is proportional and differs by a sine term.

#### Supplementary Figures

**Figure S1:**
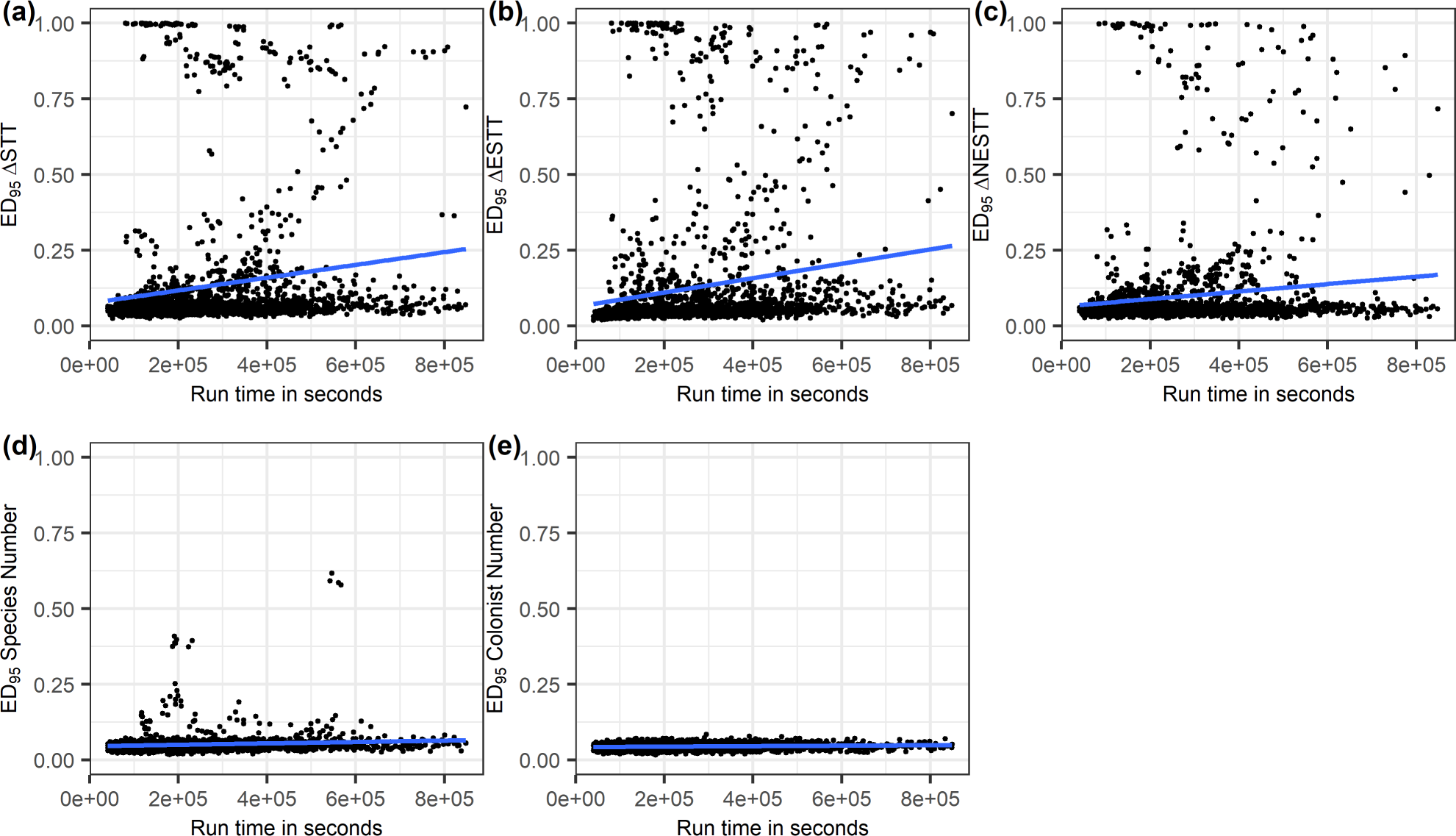
Pearson correlation between ED_95_ statistic and computational run time in the robustness pipeline (see Fig. 2, main text) for (a) ΔSTT, (b) ΔESTT, (c) ΔNESTT, (d) number of species, (e) number of colonists for all parameter spaces. Pearson’s correlation coefficient for (a) is 0.16, (b) is 0.17, (c) is 0.12, (d) is 0.11, (e) is 0.13.

**Figure S2:**
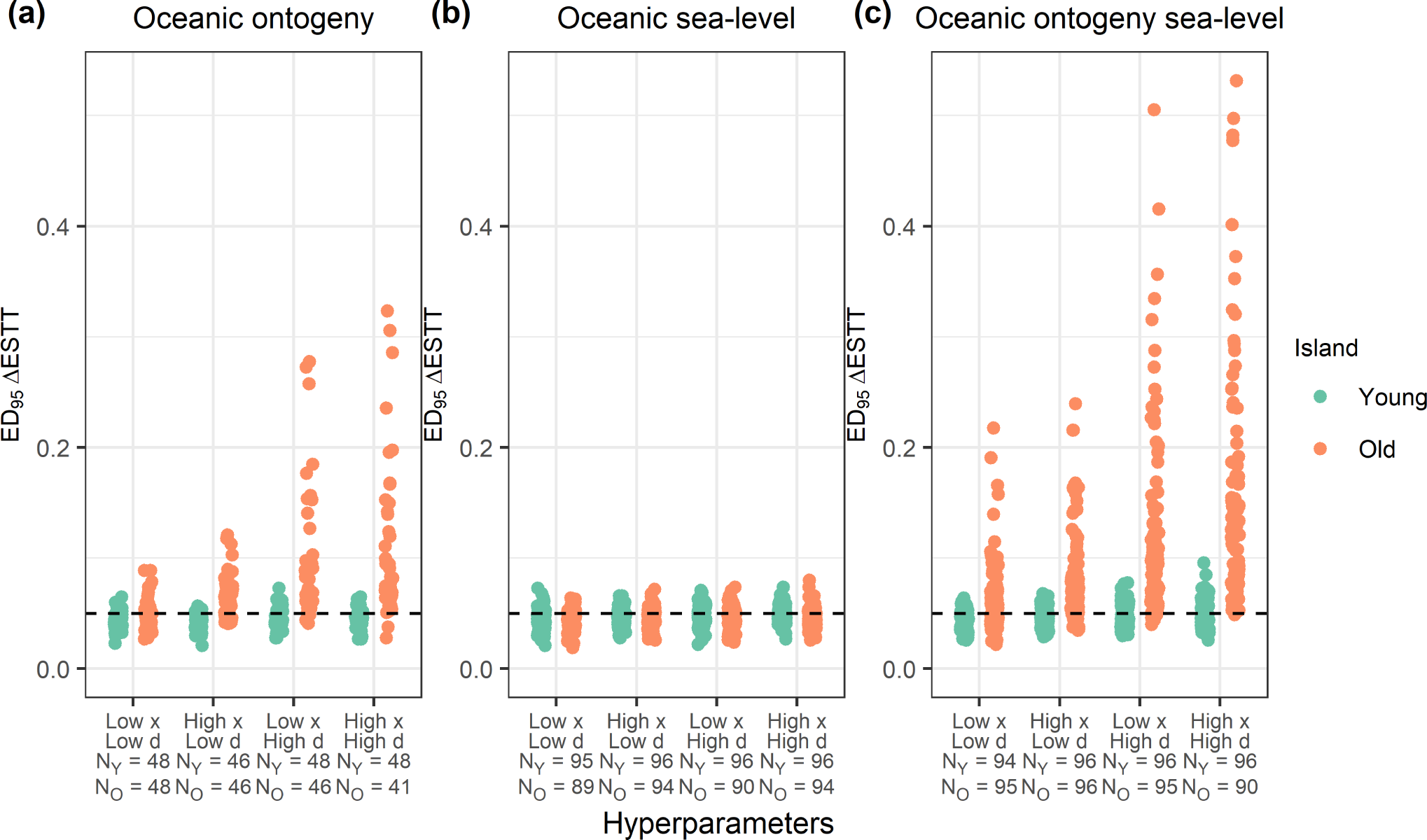
Strip charts showing the distributions of the endemic ED_95_ statistic (ΔESTT ED_95_) for each combination of hyperparameters (*d* and *x* controlling the effect of area on the rates of cladogenesis and extinction respectively). One point represents the ED_95_ for a single parameter set with the specified hyperparameters on the *x* -axis. All plots have a dashed line at 0.05 which is the null expectation of the ED_95_. (a) ΔESTT ED_95_ statistic for oceanic ontogeny. (b) ΔESTT ED_95_ statistic for oceanic sea-level. (c) ΔESTT ED_95_ statistic for oceanic ontogeny and sea-level. Sample size for young island (green) on each strip is given in the *x* -axis label by N_Y_. Sample size for old island (orange) on each strip is given in the *x* -axis label by N_O_.

**Figure S3:**
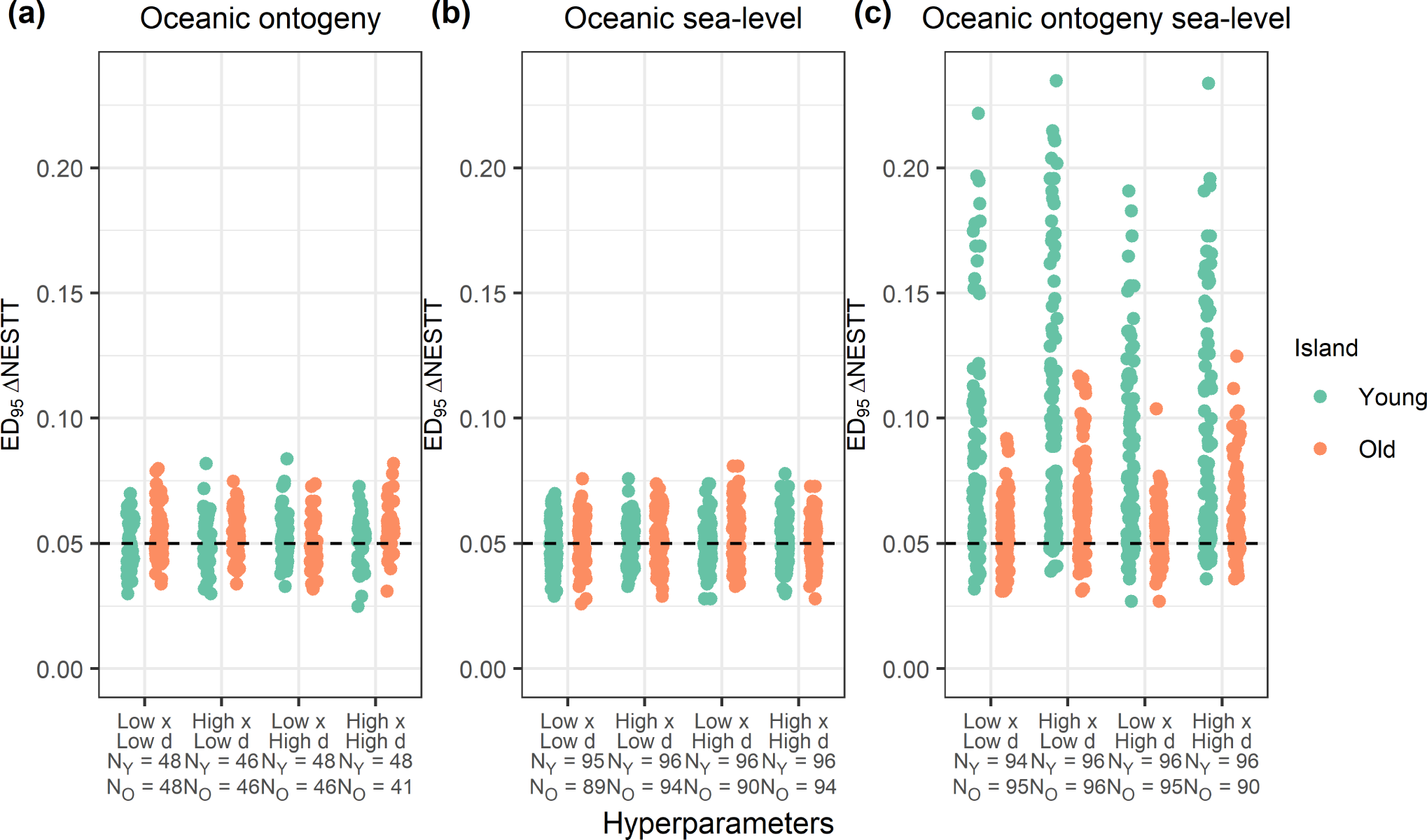
Strip charts showing the distributions of the non-endemic ED_95_ statistic (ΔNESTT ED_95_) for each combination of hyperparameters (*d* and *x* controlling the effect of area on the rates of cladogenesis and extinction respectively). One point represents the ED_95_ for a single parameter set with the specified hyperparameters on the *x* -axis. All plots have a dashed line at 0.05 which is the null expectation of the ED_95_. (a) ΔNESTT ED_95_ statistic for oceanic ontogeny. (b) ΔNESTT ED_95_ statistic for oceanic sea-level. (c) ΔNESTT ED_95_ statistic for oceanic ontogeny and sea-level. Sample size for young island (green) on each strip is given in the *x* -axis label by N_Y_. Sample size for old island (orange) on each strip is given in the *x* -axis label by N_O_.

**Figure S4:**
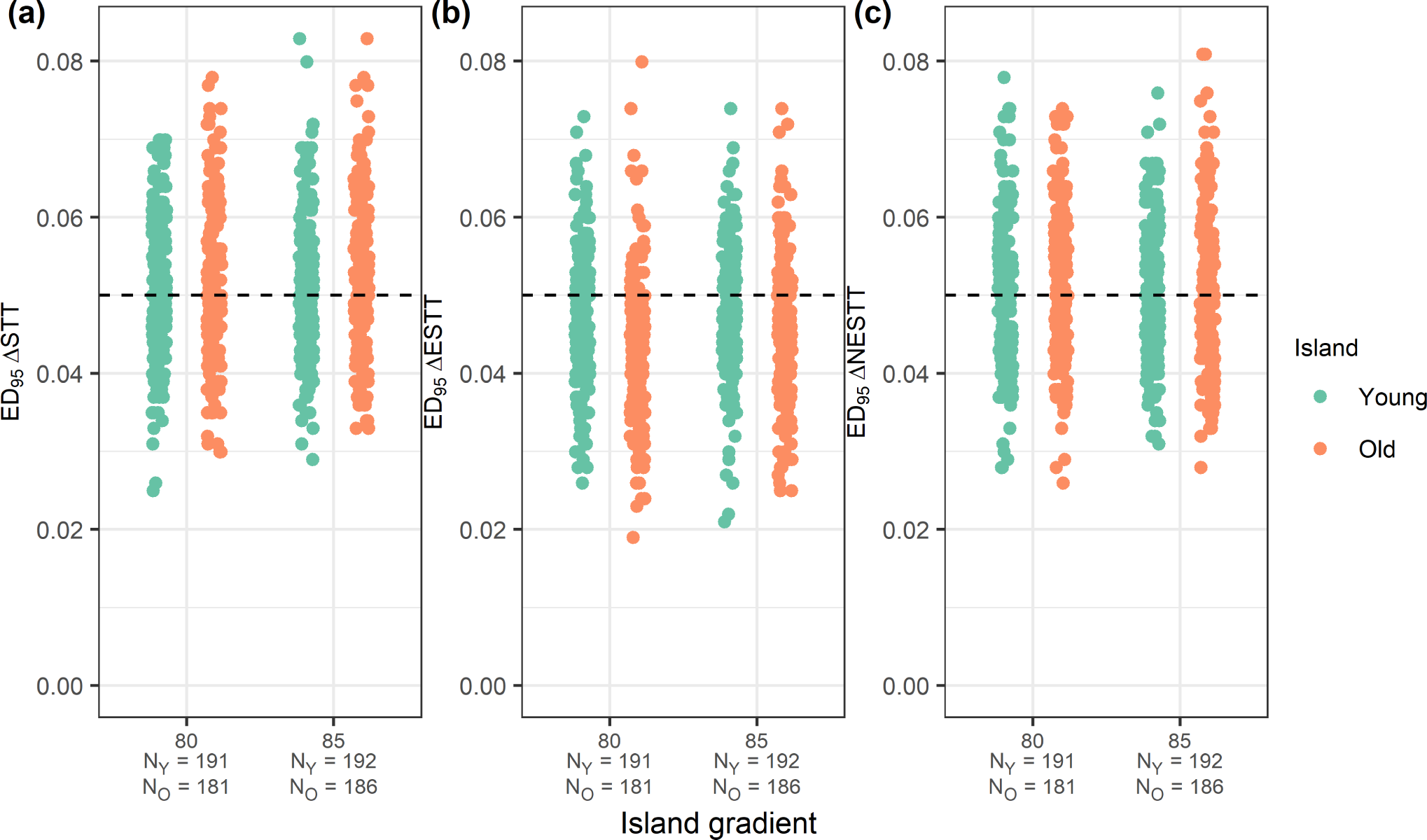
Strip charts showing the distributions of the ED_95_ statistic for each combination of island gradients for oceanic sea-level (without ontogeny). One point represents the ED_95_ for a single parameter set with the specified island gradient on the *x* -axis. All plots have a dashed line at 0.05 which is the null expectation of the ED_95_. (a) ΔSTT ED_95_ statistic, (b) ΔESTT ED_95_ statistic, (c) ΔNESTT ED_95_ statistic for oceanic sea-level. Sample size for young island (green) on each strip is given in the *x* -axis label by N_Y_. Sample size for old island (orange) on each strip is given in the *x* -axis label by N_O_.

**Figure S5:**
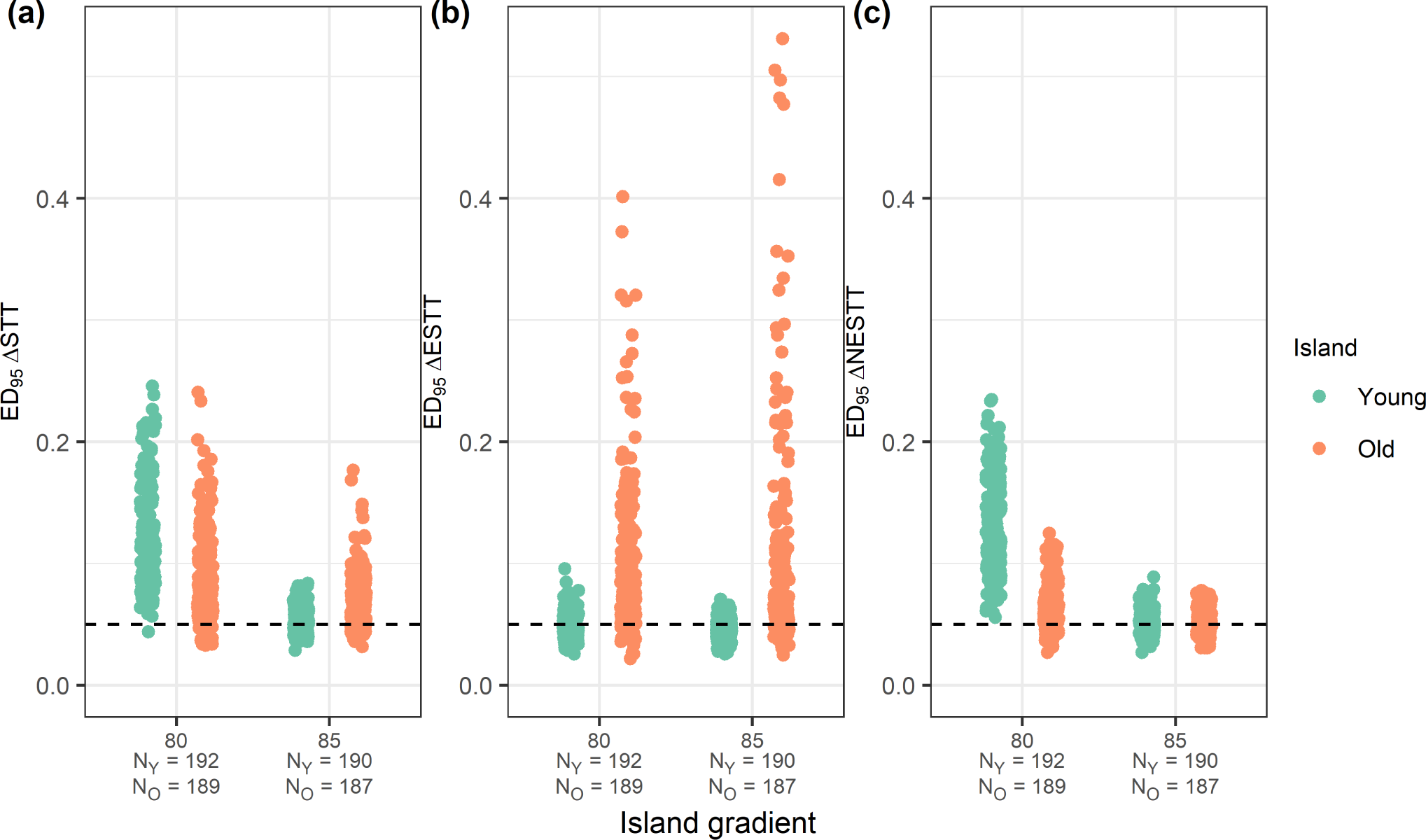
Strip charts showing the distributions of the ED_95_ statistic for each combination of island gradients for oceanic ontogeny sea-level. One point represents the ED_95_ for a single parameter set with the specified island gradient on the *x* -axis. All plots have a dashed line at 0.05 which is the null expectation of the ED_95_. (a) ΔSTT ED_95_ statistic, (b) ΔESTT ED_95_ statistic, (c) ΔNESTT ED_95_ statistic for oceanic ontogeny sea-level. Sample size for young island (green) on each strip is given in the *x* -axis label by N_Y_. Sample size for old island (orange) on each strip is given in the *x* -axis label by N_O_.

**Figure S6:**
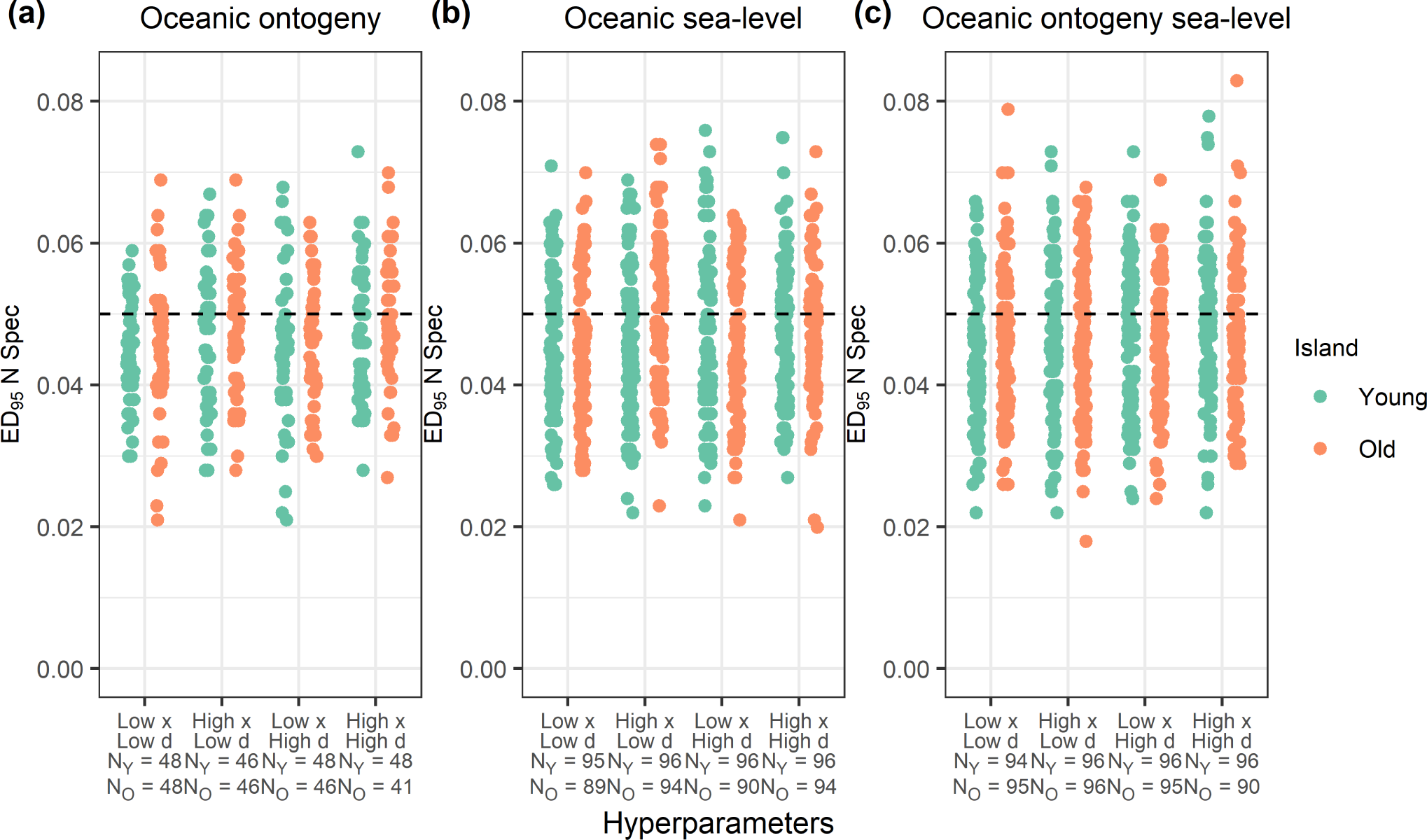
Strip charts showing the distributions of the ED_95_ statistic for number of species at the present (at the end of the simulation) for each combination of hyperparameters (*d* and *x* controlling the effect of area on the rates of cladogenesis and extinction respectively). One point represents the ED_95_ for a single parameter set with the specified hyperparameters on the *x* -axis. All plots have a dashed line at 0.05 which is the null expectation of the ED_95_. (a) ED_95_ statistic for number of species at the present for oceanic ontogeny. (b) ED_95_ statistic for number of species at the present for oceanic sea-level. (c) ED_95_ statistic for number of species at the present for oceanic ontogeny and sea-level. Sample size for young island (green) on each strip is given in the *x* -axis label by N_Y_. Sample size for old island (orange) on each strip is given in the *x* -axis label by N_O_.

**Figure S7:**
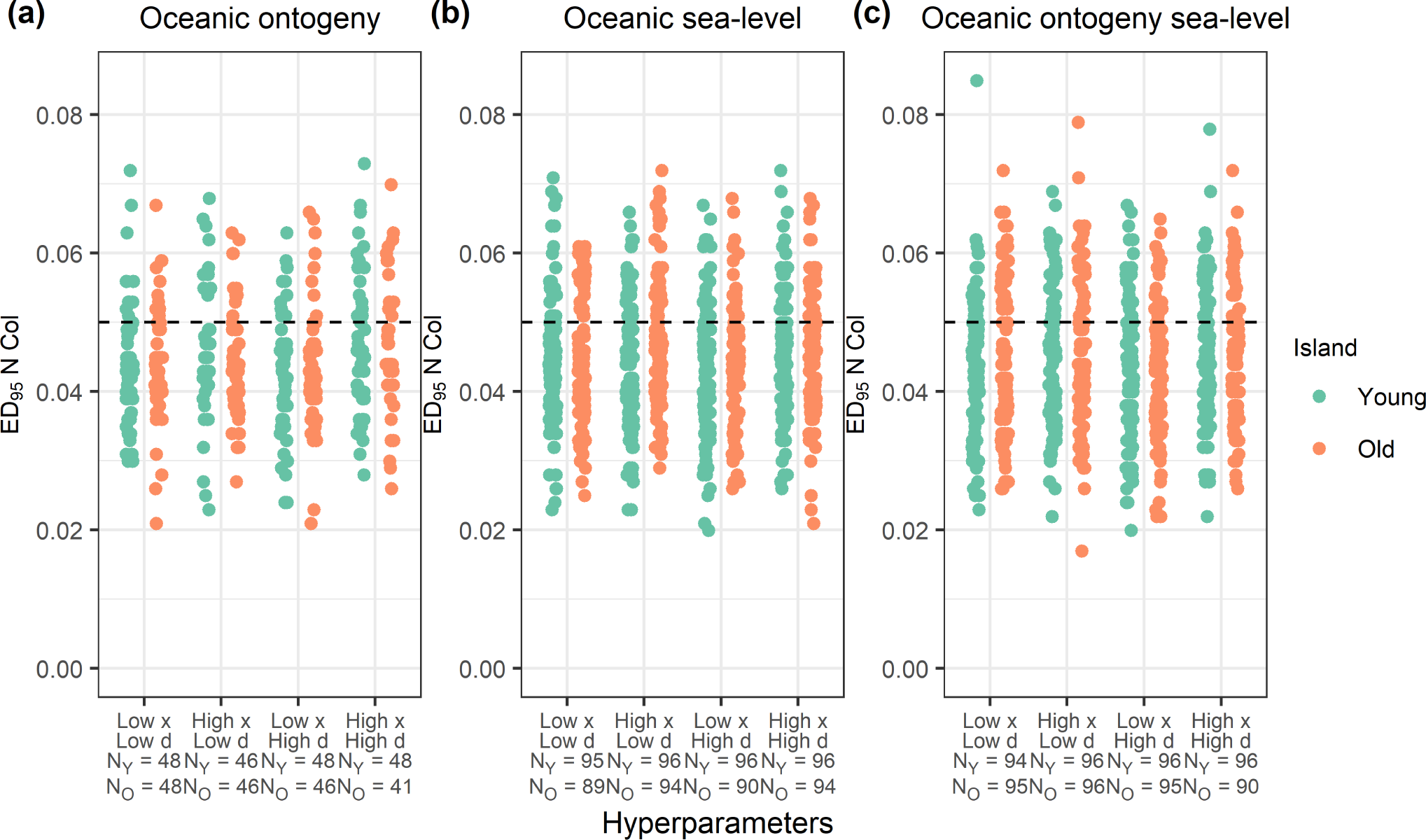
Strip charts showing the distributions of the ED_95_ statistic for number of colonists at the present (at the end of the simulation) for each combination of hyperparameters (*d* and *x* controlling the effect of area on the rates of cladogenesis and extinction respectively). One point represents the ED_95_ for a single parameter set with the specified hyperparameters on the *x* -axis. All plots have a dashed line at 0.05 which is the null expectation of the ED_95_. (a) ED_95_ statistic for number of colonists at the present for oceanic ontogeny. (b) ED_95_ statistic for number of colonists at the present for oceanic sea-level. (c) ED_95_ statistic for number of colonists at the present for oceanic ontogeny and sea-level. Sample size for young island (green) on each strip is given in the *x* -axis label by N_Y_. Sample size for old island (orange) on each strip is given in the *x* -axis label by N_O_.

**Figure S8:**
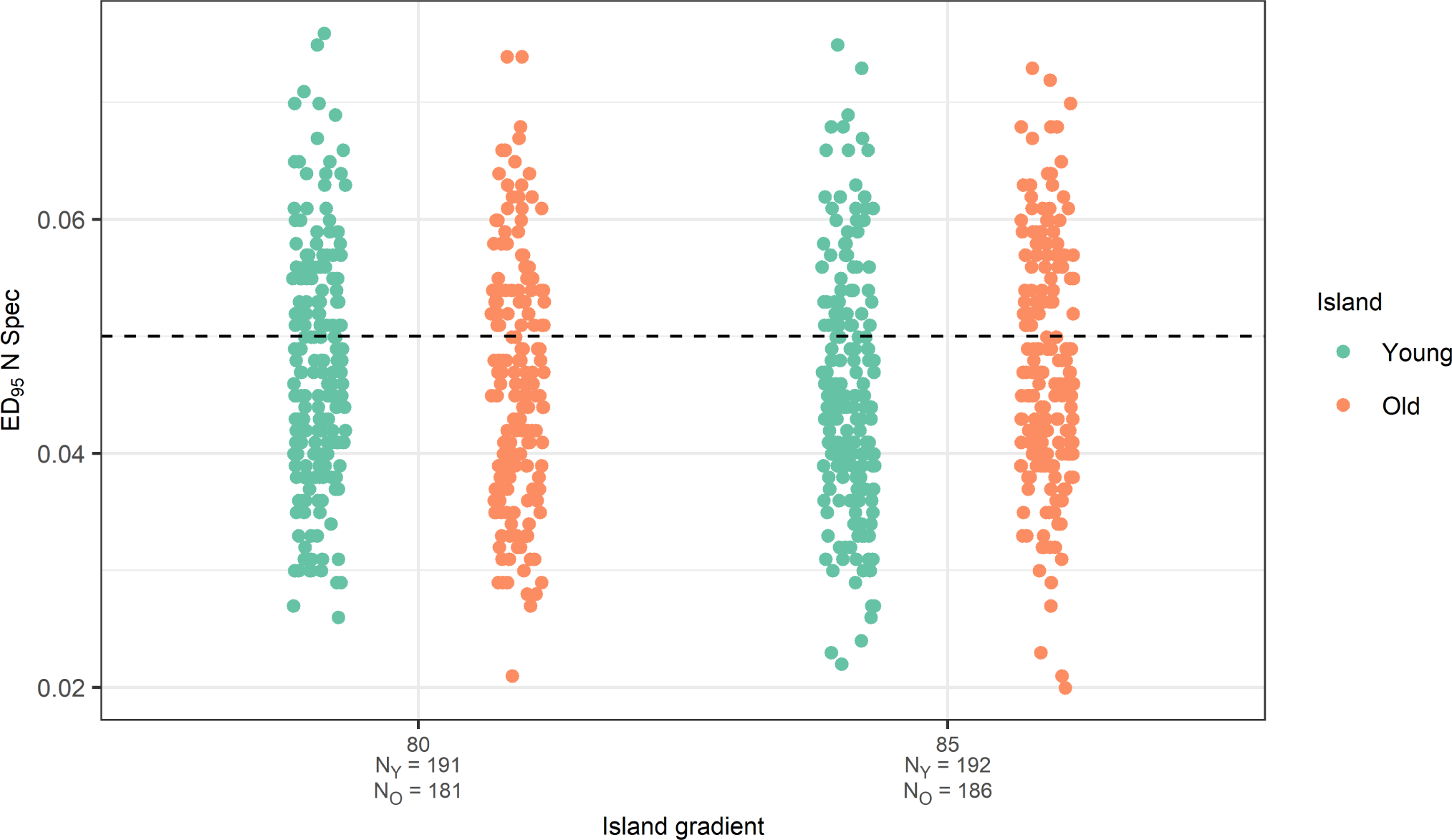
Strip charts showing the distributions of the ED_95_ statistic for the number of species at the present (N Spec) for each combination of island gradients for oceanic sea-level. One point represents the ED_95_ for a single parameter set with the specified island gradient on the *x* -axis. All plots have a dashed line at 0.05 which is the null expectation of the ED_95_. ED_95_ statistic for number of species at the present for oceanic sea-level. Sample size for young island (green) on each strip is given in the *x* -axis label by N_Y_. Sample size for old island (orange) on each strip is given in the *x* -axis label by N_O_.

**Figure S9:**
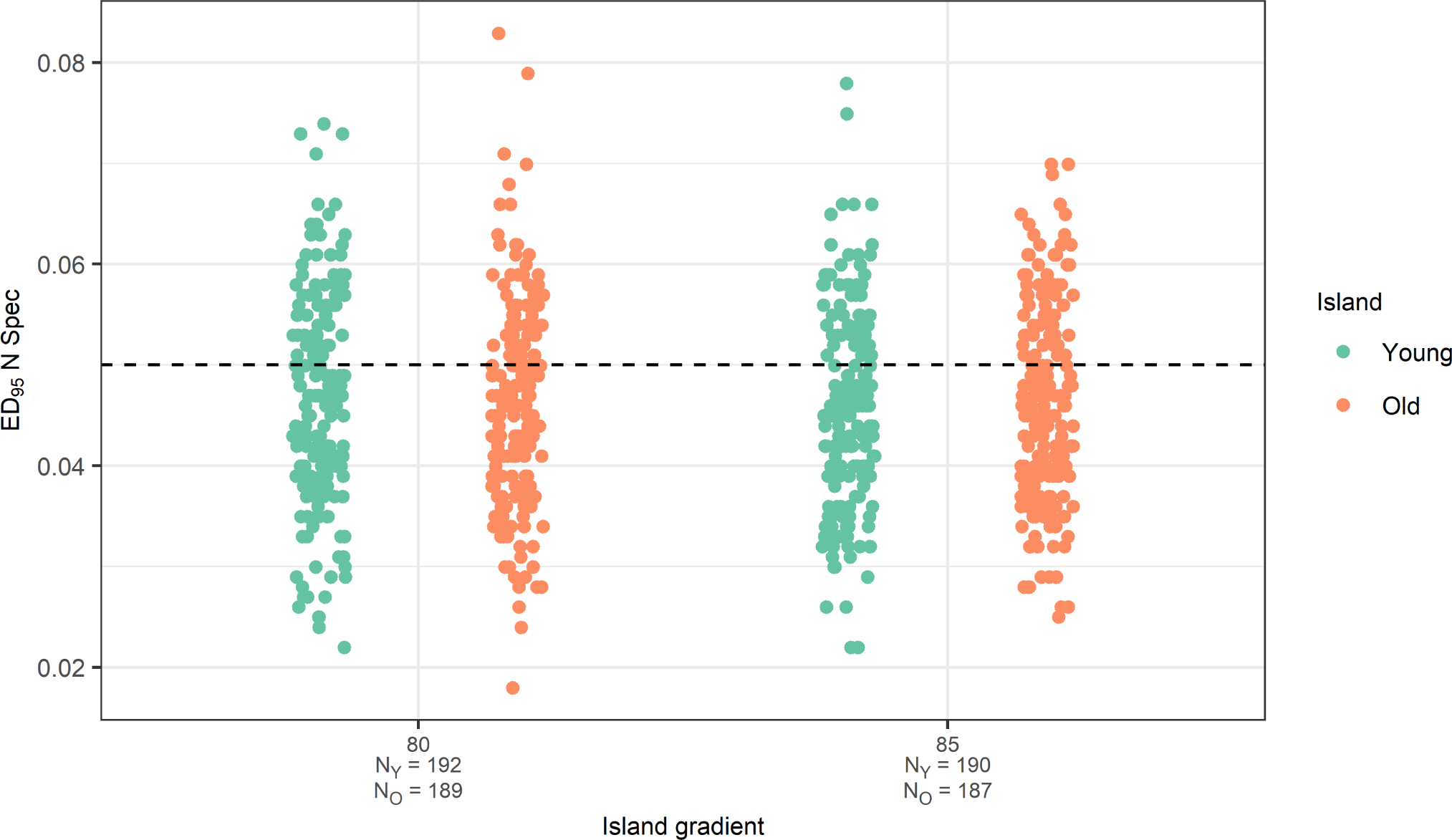
Strip charts showing the distributions of the ED_95_ statistic for the number of species at the present (N Spec) for each combination of island gradients for oceanic ontogeny sea-level. One point represents the ED_95_ for a single parameter set with the specified island gradient on the *x* -axis. All plots have a dashed line at 0.05 which is the null expectation of the ED_95_. ED_95_ statistic for number of species at the present for oceanic ontogeny sea-level. Sample size for young island (green) on each strip is given in the *x* -axis label by N_Y_. Sample size for old island (orange) on each strip is given in the *x* -axis label by N_O_.

**Figure S10:**
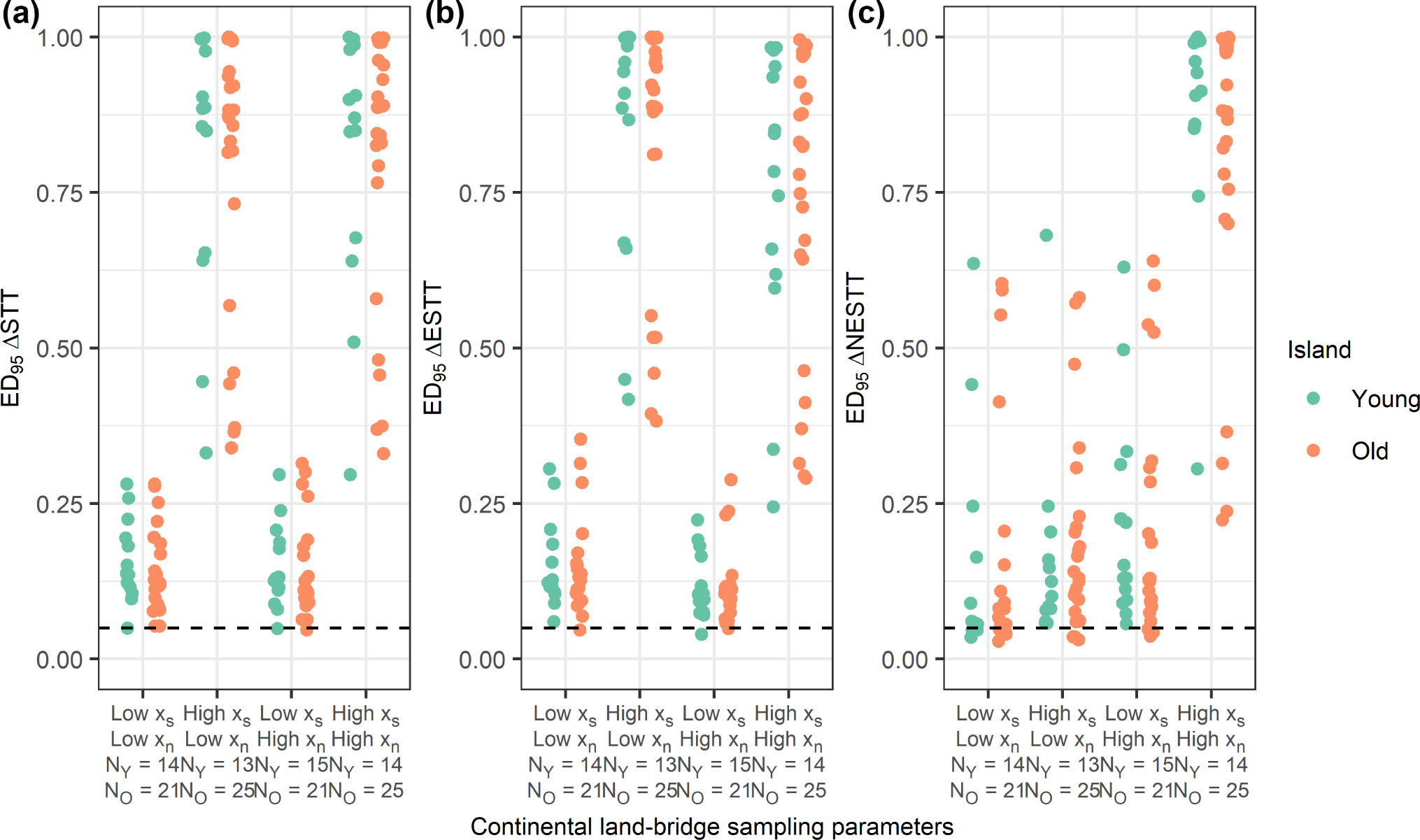
Strip charts showing the distributions of the ED_95_ statistic for each combination of continental sampling parameters (*x_s_* and *x_n_* controlling the sampling probability of a species being initially present on the island and the sampling probability of species present initially on the island being non-endemic). One point represents the ED_95_ for a single parameter set with the specified sampling parameters on the *x* -axis. All plots have a dashed line at 0.05 which is the null expectation of the ED_95_. (a) ΔSTT ED_95_ statistic, (b) ΔESTT ED_95_ statistic, (c) ΔNESTT ED_95_ statistic for continental land-bridge. Sample size for young island (green) on each strip is given in the *x* -axis label by N_Y_. Sample size for old island (orange) on each strip is given in the *x* -axis label by N_O_.

**Figure S11:**
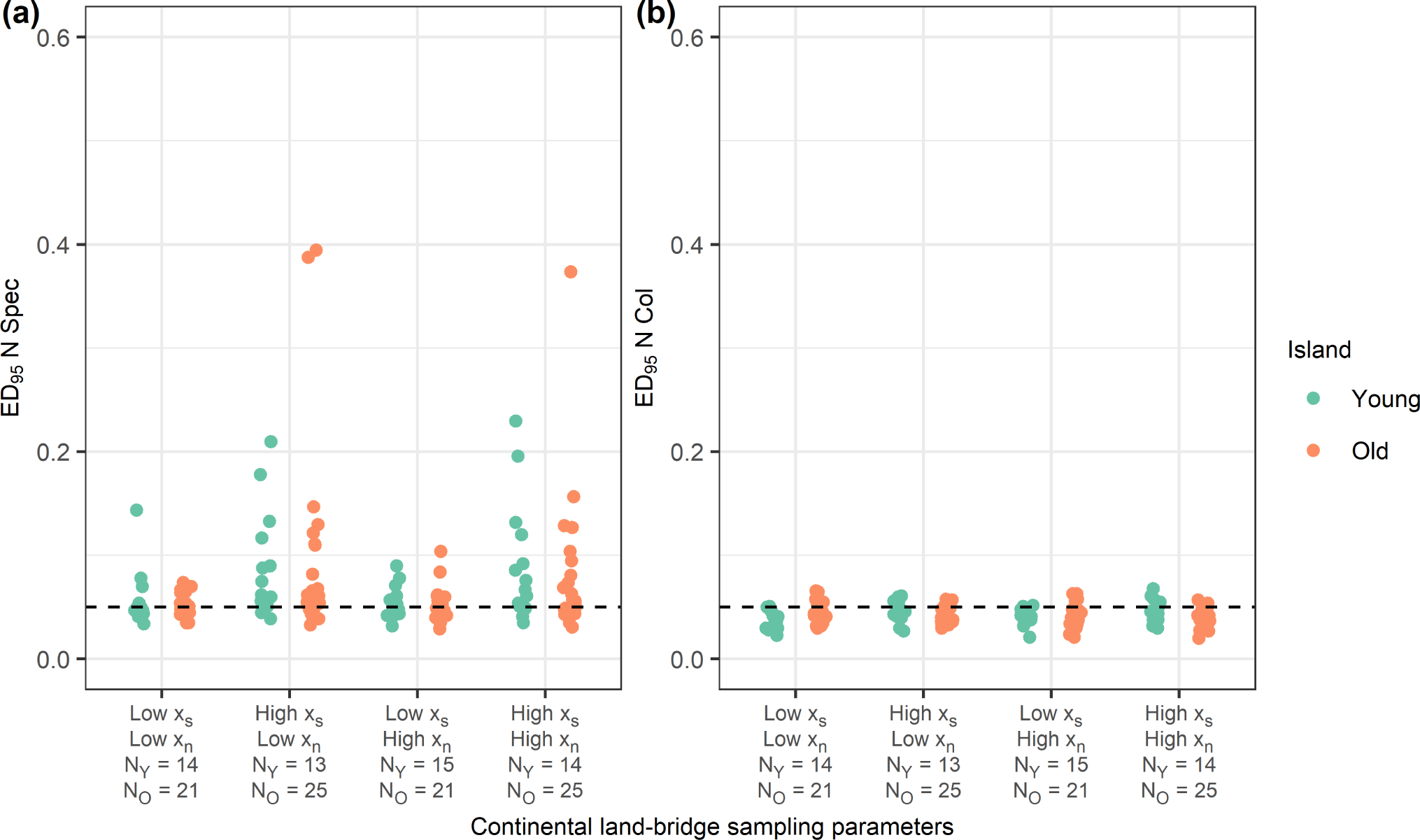
Strip charts showing the distributions of the ED_95_ statistic for number of species (N Spec) (a) and colonists (N Col) (b) at the present for each combination of continental sampling parameters (*x_s_* and *x_n_* controlling the sampling probability of a species being initially present on the island and the sampling probability of species present initially on the island being nonendemic). One point represents the ED_95_ for a single parameter set with the specified sampling parameters on the *x* -axis. All plots have a dashed line at 0.05 which is the null expectation of the ED_95_. (a) ΔSTT ED_95_ statistic, (b) ΔESTT ED_95_ statistic, (c) ΔNESTT ED_95_ statistic for continental land-bridge. Sample size for young island (green) on each strip is given in the *x* -axis label by N_Y_. Sample size for old island (orange) on each strip is given in the *x* -axis label by N_O_.

**Figure S12:**
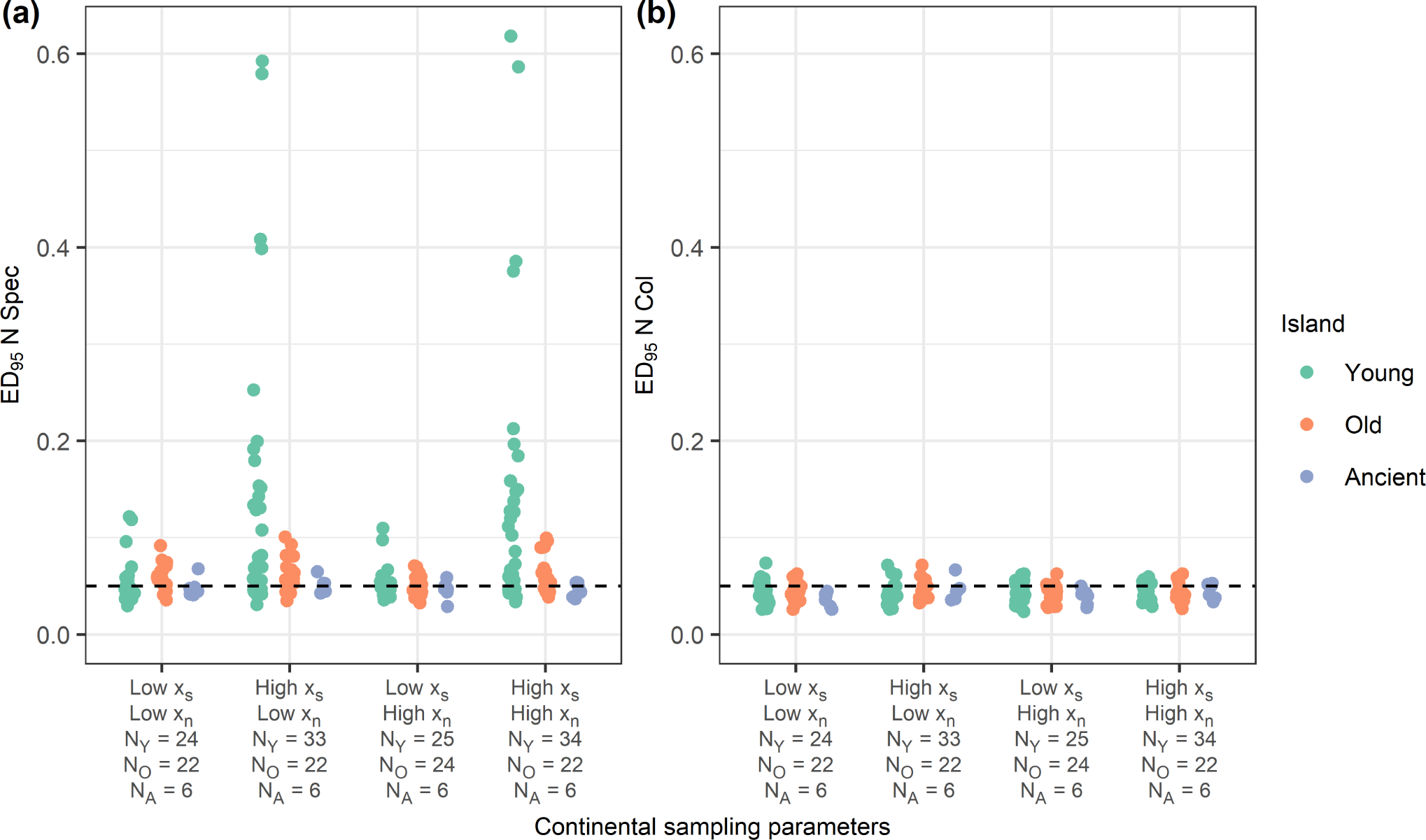
Strip charts showing the distributions of the ED_95_ statistic for number of species and colonists at the present for each combination of continental sampling parameters (*x_s_* and *x_n_* controlling the sampling probability of a species being initially present on the island and the sampling probability of species present initially on the island being non-endemic). One point represents the ED_95_ for a single parameter set with the specified sampling parameters on the *x* - axis. All plots have a dashed line at 0.05 which is the null expectation of the ED_95_. Sample size for young island (green) on each strip is given in the *x* -axis label by N_Y_. Sample size for old island (orange) on each strip is given in the *x* -axis label by N_O_. Sample size for the ancient island (blue) on each strip is given in the *x* -axis label by N_A_.

#### Supplementary Tables

**Table S1:** Parameter space for oceanic ontogeny young island. Parameter space consists of each combination of the model parameters.

| **Oceanic ontogeny young island** |  |
| --- | --- |
| **Model parameters** | **Parameter value** |
| Time | 2.55 |
| Mainland species pool | 1000 |
| Cladogenesis | 0.02, 0.04 |
| Extinction | 0.975, 1.95 |
| Carrying capacity | 0.001, 0.01, ∞ |
| Colonisation | 0.03363, 0.06726 |
| Anagenesis | 0.0295, 0.059 |
| $x$ | 0.075, 0.15 |
| $d$ | 0.1108, 0.2216 |
| Max area | 13500 |
| Current area | 3155 |
| Peak time | 0.53 |
| Total island age | 2.864 |

**Table S2:** Parameter space for oceanic ontogeny old island. Parameter space consists of each combination of the model parameters.

| Oceanic ontogeny old island |  |
| --- | --- |
| Model parameters | Parameter value |
| Time | 6.15 |
| Mainland species pool | 1000 |
| Cladogenesis | 0.02, 0.04 |
| Extinction | 0.975, 1.95 |
| Carrying capacity | 0.001, 0.01, ∞ |
| Colonisation | 0.03363, 0.06726 |
| Anagenesis | 0.0295, 0.059 |
| $x$ | 0.075, 0.15 |
| $d$ | 0.1108, 0.2216 |
| Max area | 3787 |
| Current area | 1431 |
| Peak time | 0.27 |
| Total island age | 8.473 |

**Table S3:** Parameter space for oceanic sea-level young island. Parameter space consists of each combination of the model parameters.

| Oceanic sea-level young island |  |
| --- | --- |
| Model parameters | Parameter value |
| Time | 2.55 |
| Mainland species pool | 1000 |
| Cladogenesis | 0.02, 0.04 |
| Extinction | 0.975, 1.95 |
| Carrying capacity | 0.001, 0.01, ∞ |
| Colonisation | 0.03363, 0.06726 |
| Anagenesis | 0.0295, 0.059 |
| $x$ | 0.075, 0.15 |
| $d$ | 0.1108, 0.2216 |
| Current area | 3155 |
| Sea-level amplitude | 60 |
| Sea-level frequency | 25.5 |
| Island gradient angle | 80, 85 |

**Table S4:** Parameter space for oceanic sea-level old island. Parameter space consists of each combination of the model parameters.

| Oceanic sea-level old island |  |
| --- | --- |
| Model parameters | Parameter value |
| Time | 6.15 |
| Mainland species pool | 1000 |
| Cladogenesis | 0.02, 0.04 |
| Extinction | 0.975, 1.95 |
| Carrying capacity | 0.001, 0.01, ∞ |
| Colonisation | 0.03363, 0.06726 |
| Anagenesis | 0.0295, 0.059 |
| $x$ | 0.075, 0.15 |
| $d$ | 0.1108, 0.2216 |
| Current area | 1431 |
| Sea-level amplitude | 60 |
| Sea-level frequency | 61.5 |
| Island gradient angle | 80, 85 |

**Table S5:** Parameter space for oceanic ontogeny sea-level young island. Parameter space consists of each combination of the model parameters.

| **Oceanic ontogeny sea-level young island** |  |
| --- | --- |
| **Model parameters** | **Parameter value** |
| Time | 2.55 |
| Mainland species pool | 1000 |
| Cladogenesis | 0.02, 0.04 |
| Extinction | 0.975, 1.95 |
| Carrying capacity | 0.001, 0.01, ∞ |
| Colonisation | 0.03363, 0.06726 |
| Anagenesis | 0.0295, 0.059 |
| $x$ | 0.075, 0.15 |
| $d$ | 0.1108, 0.2216 |
| Max area | 13500 |
| Current area | 3155 |
| Peak time | 0.53 |
| Total island age | 2.864 |
| Sea-level amplitude | 60 |
| Sea-level frequency | 25.5 |
| Island gradient angle | 80, 85 |

**Table S6:** Parameter space for oceanic ontogeny sea-level old island. Parameter space consists of each combination of the model parameters.

| **Oceanic ontogeny sea-level old island** |  |
| --- | --- |
| **Model parameters** | **Parameter value** |
| Time | 6.15 |
| Mainland species pool | 1000 |
| Cladogenesis | 0.02, 0.04 |
| Extinction | 0.975, 1.95 |
| Carrying capacity | 0.001, 0.01, ∞ |
| Colonisation | 0.03363, 0.06726 |
| Anagenesis | 0.0295, 0.059 |
| $x$ | 0.075, 0.15 |
| $d$ | 0.1108, 0.2216 |
| Max area | 3787 |
| Current area | 1431 |
| Peak time | 0.27 |
| Total island age | 8.473 |
| Sea-level amplitude | 60 |
| Sea-level frequency | 61.5 |
| Island gradient angle | 80, 85 |

**Table S7:** Parameter space for continental young, old and ancient islands. Parameter space consists of each combination of the model parameters.

| **Continental young, old, and ancient islands** |  |
| --- | --- |
| **Model parameters** | **Parameter value** |
| Time | 2.55, 6.15, 50.00 |
| Mainland species pool | 1000 |
| Cladogenesis | 0.5, 1.0 |
| Extinction | 0.5, 1.0 |
| Carrying capacity | 10, 40, ∞ |
| Colonisation | 0.01, 0.05 |
| Anagenesis | 1.0, 2.0 |
| $x_s$ | 0.01, 0.05 |
| $x_n$ | 0.1, 0.9 |

**Table S8:**
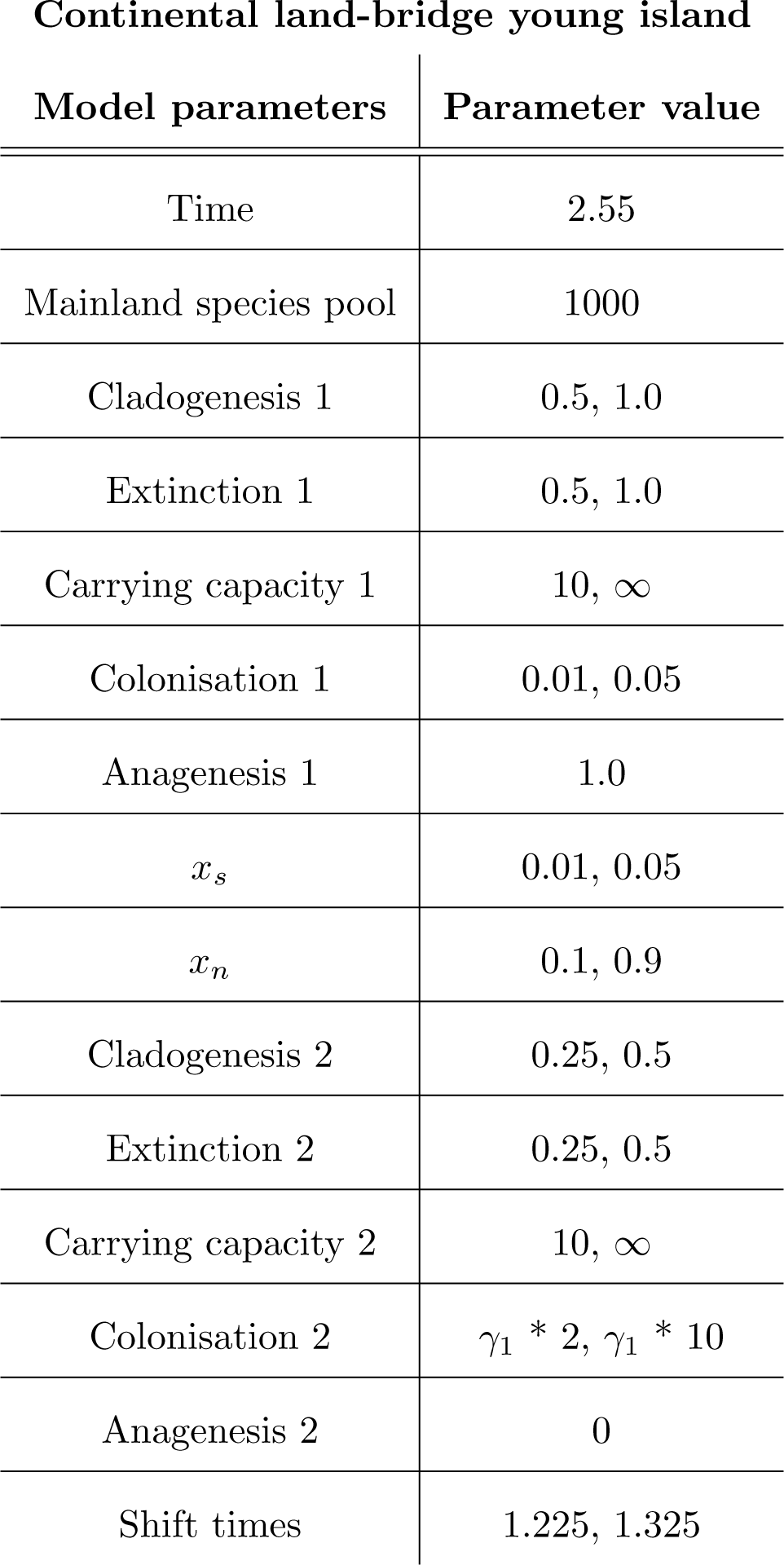
Parameter space for continental land-bridge young island. Parameter space consists of each combination of the model parameters. Colonisation rate 2 (i.e. when the land-bridge is present) is colonisation rate 1 (*γ*_1_), multipled by a colonisation rate multiplier, which in our case is two and ten.

| **Continental land-bridge young island** |  |
| --- | --- |
| **Model parameters** | **Parameter value** |
| Time | 2.55 |
| Mainland species pool | 1000 |
| Cladogenesis 1 | 0.5, 1.0 |
| Extinction 1 | 0.5, 1.0 |
| Carrying capacity 1 | 10, ∞ |
| Colonisation 1 | 0.01, 0.05 |
| Anagenesis 1 | 1.0 |
| $x_s$ | 0.01, 0.05 |
| $x_n$ | 0.1, 0.9 |
| Cladogenesis 2 | 0.25, 0.5 |
| Extinction 2 | 0.25, 0.5 |
| Carrying capacity 2 | 10, ∞ |
| Colonisation 2 | $\gamma_1 * 2, \gamma_1 * 10$ |
| Anagenesis 2 | 0 |
| Shift times | 1.225, 1.325 |

**Table S9:** Parameter space for continental land-bridge old island. Parameter space consists of each combination of the model parameters. Colonisation rate 2 (i.e. when the land-bridge is present) is colonisation rate 1 (*γ*_1_), multipled by a colonisation rate multiplier, which in our case is two and ten.

| **Continental land-bridge old island** |  |
| --- | --- |
| **Model parameters** | **Parameter value** |
| Time | 6.15 |
| Mainland species pool | 1000 |
| Cladogenesis 1 | 0.5, 1.0 |
| Extinction 1 | 0.5, 1.0 |
| Carrying capacity 1 | 10, ∞ |
| Colonisation 1 | 0.01, 0.05 |
| Anagenesis 1 | 1.0 |
| $x_s$ | 0.01, 0.05 |
| $x_n$ | 0.1, 0.9 |
| Cladogenesis 2 | 0.25, 0.5 |
| Extinction 2 | 0.25, 0.5 |
| Carrying capacity 2 | 10, ∞ |
| Colonisation 2 | $\gamma_1 * 2, \gamma_1 * 10$ |
| Anagenesis 2 | 0 |
| Shift times | 3.025, 3.125 |

